# Molecular landscape of BoNT/B bound to a membrane-inserted synaptotagmin/ganglioside complex

**DOI:** 10.1101/2021.10.18.464377

**Authors:** Jorge Ramirez-Franco, Fodil Azzaz, Marion Sangiardi, Géraldine Ferracci, Fahamoe Youssouf, Michel Robert Popoff, Michael Seagar, Christian Lévêque, Jacques Fantini, Oussama El Far

**Author notes:** These authors contributed equally to this work (Jorge Ramirez-Franco, Fodil Azzaz).

## Abstract

Botulinum neurotoxin serotype B (BoNT/B) uses two separate protein and polysialoglycolipid-binding pockets to interact with synaptotagmin 1/2 and gangliosides. However, an integrated model of BoNT/B bound to its neuronal receptors in a native membrane topology is still lacking. Using a panel of in silico and experimental approaches, we present here a new model for BoNT/B binding to neuronal membranes, in which the toxin binds to a preassembled synaptotagmin-ganglioside GT1b complex and a free ganglioside allowing a lipid-binding loop of BoNT/B to interact with the glycone part of the synaptotagmin-associated GT1b. Furthermore, our data provide molecular support for the decrease in BoNT/B sensitivity in Felidae that harbor the natural variant synaptotagmin2-N_59_Q. These results reveal multiple interactions of BoNT/B with gangliosides and support a novel paradigm in which a toxin recognizes a protein/ganglioside complex.

## Introduction

Botulinum neurotoxins (BoNTs), are a family of potent protein toxins produced by anaerobic gram-positive Clostridia ^1 2^. BoNTs are classified as seven different serotypes from BoNT/A to G, further divided into subtypes with different amino acid sequences, although additional BoNTs are still being discovered, including mosaic toxins derived from a combination of different serotypes ^1 3^.

BoNTs are the etiological agents of botulism, a rare but severe disease affecting many vertebrates, which results from the inhibition of acetylcholine release in the peripheral nervous system, causing flaccid paralysis. At the same time, BoNT/A, and to a lesser extent BoNT/B, are widely exploited for therapeutic applications ^4 5 6^.

BoNTs intoxicate neurons using a multistep mechanism. BoNTs are structurally similar to AB toxins with a 100 kDa heavy chain (HC) and a 50 kDa catalytic light chain, associated via a disulphide bond and non-covalent interactions. After entering the circulation, BoNTs target high affinity receptors on peripheral nerve terminals via the HC domain. The amino-terminal domain of the HC then translocates the enzymatic light chain into the cytoplasm where the latter cleaves one of the three intracellular SNARE proteins (VAMP1-3, SNAP-25 or syntaxin 1) necessary for synaptic vesicle fusion and neurotransmitter release ^5^. The BoNT carboxyl-terminal sub-domain of HC, organized into a β-trefoil fold, displays two independent binding pockets, which bind two distinct classes of receptors.

The first receptor to be discovered is a ganglioside ^7^, typically GT1b or GD1a, that is recognized by a ganglioside-binding site (GBS1), conserved in most BoNTs. Gangliosides associate with cholesterol in tightly packed lipid domains that are in dynamic equilibrium with less ordered membrane regions and can support lipid and protein-lipid interactions in *cis* and *trans* configurations ^7 8^. Moreover, microdomains containing gangliosides provide an entry pathway for several viruses and other pathogens ^9 10 11^.

The second receptor is a protein, corresponding to the luminal sequence of a synaptic vesicle protein: synaptotagmin 1 and 2 (SYT) for BoNT/B, G and the mosaic toxin BoNT/DC or SV2 for BoNT/A, D, E, F although the identity of BoNT/D protein receptor is still to be confirmed ^2^. BoNT/C has no identified protein receptor, but like BoNT/D, harbors an additional ganglioside-binding pocket distinct from GBS1 and termed “sialic-binding site” that overlaps with the SYT-binding pocket of BoNT/B, contributing to toxicity ^12 13^.

Besides these two receptors, a solvent-exposed lipid-binding loop (LBL), present in BoNT/B, C, D, G and BoNT/DC is localized between GBS1 and the protein (BoNT/B, G and DC) or sialic acid-binding pocket (BoNT/C and BoNT/D), participates in BoNT toxicity and neuronal membranes recognition ^2 14 15^. The exceptional neurotropism of BoNT/B is conferred by interaction with the extracellular juxtamembrane domain (JMD) of SYT, that is translocated to the plasma membrane by synaptic vesicle fusion. SYT1 and SYT2 have comparable biochemical properties and similar functions, regulating exo-endocytic recycling of synaptic vesicles by interacting with cytosolic proteins such as the adaptor protein AP2, as well as with specific lipids like cholesterol and PIP2 ^16 17 18^. While SYT1 is widely distributed in terminals of autonomic and sensory neurons, as well as in some neuromuscular junctions, SYT2 is the dominant isoform at most neuromuscular junctions ^5^. Co-crystallization data indicate that BoNT/B binds SYT1 and SYT2 in a very similar manner using a saddle-shaped pocket interacting with 10– 14 SYT JMD residues ^19 20 21 6^. The extracellular domain of SYT is not structured in solution, but the JMD of SYT adopts a helical conformation upon binding to BoNT/B ^19 20 21 6^. In the absence of gangliosides, BoNT/B displays a much higher affinity for rat SYT2 (40 nM) than for SYT1, due to a small difference in primary sequence in the SYT JMD ^19^. Although BoNT/B has low affinity for GT1b (µM range) ^14 15^, the latter drastically increases BoNT/B affinity (∼0.4 nM) for membranes containing SYT ^22 23 24^. In detergent, the synergistic effect of GT1b requires the presence of the SYT transmembrane domain (TMD) ^25 26^ and high affinity binding is only reached in reconstituted lipids systems containing GT1b as well as the transmembrane domain of SYT, suggesting a role for the intramembrane segments in toxin binding ^21 23^. As the available structural data were obtained in the absence of apolar domains of BoNT/B receptors ^20 19 21 6^, how BoNT/B binds to its receptors in a membrane context remains to be elucidated.

Recently we reported that the transmembrane and juxtamembrane domains of SYT interact with complex gangliosides inducing an α–helical structure ^24^. A mutation (SYT1-K52A) that decreases GT1b assembly with SYT1, abolished BoNT/B binding in neuroendocrine cells, suggesting that the preassembly of a GT1b/SYT complex is crucial for BoNT/B interaction. Using a panel of in silico and experimental approaches, we now report that the SYT-binding pocket of BoNT/B can accommodate the preassembled GT1b/SYT complex. We thus propose a new model for BoNT/B-SYT interaction taking into account the membrane topology of neuronal toxin receptors, a parameter that has not been considered in previous structural studies.

## Materials and Methods

### Experimental design

The main objective of this study was to investigate how botulinum neurotoxin serotype B binds to its receptors in a membrane context, since the available structural data were obtained in the absence of apolar domains. We developed an SPR (Surface Plasmon Resonance) experimental configuration to ensure that the synaptotagmin domain that interacts with BoNT/B was bound to GT1b, before characterizing the interaction of BoNT/B with this preassembled complex. Molecular modelling was performed to model BoNT/B in interaction with the preassembled SYT1/GT1b and SYT2/GT1b complexes docked to the synaptotagmin binding pocket. For this purpose, we used BoNT/B1 structural coordinates stored in the PDB files 6G5K, 4KBB and 2NM1 to generate a complete model of BoNT/B-SYT1/2-gangliosides and cholesterol. We compared the interaction energies and landscapes of all components of the complexes with previous structural data. Newly identified contact points were compared with published reports and the importance of new synaptotagmin-contact points were experimentally validated in heterologous expression systems.

### Reagents

BoNT/B (B1 Okra strain) with non-toxic accessory proteins was from Metabiologics (Madison, WI). All peptides were synthesized by Genecust. DMPC (1,2-Dimyristoyl-sn-glycero-3-phosphocholine) was from Avanti Polar Lipids. GT1b was from Matreya LLC. Polyclonal anti-SYT1 31-55 region antibodies were generously provided by M. Takahashi. Rabbit anti-SYT2 40-65 polyclonal antibodies were produced by Genecust using a synthetic peptide (rat SYT2 40-65) and purified using protein-A sepharose. All experiments were performed in accordance with French and European guidelines for handling botulinum neurotoxin. GT1b, lyso-lactosylceramide and sphingomyelin were from Matreya LLC. Anti-BoNT/B and anti-SYT1/2 (1D12) antibodies were obtained as described ^24^. Alexa-coupled secondary antibodies were from Jackson Immunoresearch. DAPI was from SIGMA-Aldrich. Anti-GT1b monoclonal antibodies were from Merck Millipore (MAB 5608).

### SPR experiments

SPR measurements were performed with a Biacore T200 apparatus (Cytiva) using HBS (10 mM HEPES/NaOH pH 7.4, 150 mM NaCl) or TBS (10 mM TRIS/NaOH pH 7.4, 150 mM NaCl) as running buffer. CMDP chips (Xantec Bioanalytics, Germany) were functionalized with neutravidin (Pierce) according to standard protocols. pSYT peptides were injected onto neutravidin to reach between 250-500 RU, depending on the experiment. GT1b (2mg/ml in methanol) was diluted contemporaneously in running buffer and injected onto pSYT sensorchips at flow rates from 5 to 40 µl/min, depending on experiments. Gangliosides were stripped from pSYT using TBS containing CHAPS 1% (8 s at 40 µl/min). The binding stoichiometry of GT1b/pSYT was calculated using the RUmax value, determined experimentally by saturating pSYT with GT1b diluted under the CMC (10 µM) ^27^. Stoichiometry = RUmax x MW of GT1b / RU pSYT x MW of pSYT). BoNT/B was injected at 30 nM at a flow rate of 5 µl/min. For GT1b potentiation of BoNT/B binding to pSYT1, GT1b was not stripped and accumulated on pSYT1 after each BoNT/B injection. Anti-GT1b antibodies (ascites, dilution x 1000) were injected for 2 min at 10 µl/min over pSYT1 and pSYT1/GT1b complex (250 RU pSYT1 ± GT1b (30-100 RU)), before and after BoNT/B interaction (20 nM for 3 min). Unless stated, non-specific signals on control flow cells (immobilized pSYT9 or activated / deactivated empty flow cell) were automatically subtracted from measurements on experimental flow cells. Due to the low binding affinity of BoNT/B to pSYT in the absence of gangliosides, a large amount of pSYT (250-500 RU) had to immobilized. Under these conditions, accurate interaction kinetics could not be measured. Measurement of pSYT2 peptide binding to GT1b-containing liposomes was performed using hydrophobic L1 sensor chips as described ^24^. Liposomes containing 100 % DMPC (control flow cell) or 92 % DMPC, 8% GT1b (experimental flow cell) were immobilized and SYT2 peptides binding measured 5 s before the end of the injection.

### Langmuir monolayers experiments

Surface pressure measurements revealing peptide-lipid interactions at the air-water interface were studied by the Langmuir film balance technique with a fully automated microtensiometer (µTROUGH SX, Kibron Inc. Helsinki, Finland) as described previously, using an initial pressure of 17.5 mN/m ^28 29 24^.

### Immunofluorescence

HEK293 or PC12 cells, were cultured on poly-L-Lysine (10 μg/ml) treated coverslips (300,000 cells per well) in DMEM containing 5% FBS, 5% HS and 1% penicillin/streptomycin mixture (complete medium). Cells were transfected with the corresponding plasmids (pIRES-EGFP-SYT2; pIRES-EGFP-K_60_A-SYT2; pIRES-EGFP-N_59_Q-SYT2; pIRES-EGFP-SYT1 or pIRES-EGFP-H_51_G-SYT1) using Lipofectamine 2000 and according to the manufacturer’s procedure. 40 hours after transfection, GT1b (10 µg/ml) was added to the wells in DMEM and incubated for 1.5 h at 37°C followed by one washing step and transfer to complete medium. BoNT/B (10 nM and 1 nM for SYT1 and SYT2 conditions respectively) was added afterwards and incubated for 30-45 min at 37°C. After a first wash with the culture medium, additional washes were performed with PBS and cells were fixed in the dark at 4°C in 4% paraformaldehyde/PBS for 15 min followed by NH_4_Cl washing steps. Non-specific binding was blocked with 0.2% (w/v) gelatine or 5% (v/v) goat serum in a PBS buffer containing 0.1% saponin. Anti-BoNT/B (0.5 µg/µl), and 1D12 anti-SYT (1 µg/ml) antibodies were then added for 45 min at 22°C. After subsequent washing, staining was visualized using secondary anti-rabbit Alexa-594 and anti-mouse Alexa-488 antibodies. Nuclei were detected using DAPI.

### Image acquisition and analysis

Confocal images were acquired on a Zeiss LSM780 microscope and processed using ImageJ (http://rsb.info.nih.gov/ij/). For quantification, SYT immunolabeling images were thresholded in order to get a binary mask. This binary mask was used to obtain immunoreactivity (IR) values of the regions of interest (ROIs) over SYT and BoNT/B channels. For comparisons of BoNT/B binding to WT vs mutant SYTs, IR values were normalized to WT in every experiment. Results are presented as mean ± SEM. Statistical analysis was performed using Mann-Whitney U test.

### Molecular modelling

Molecular modelling studies were performed in vacuo using Hyperchem (http://www.hyper.com), Deep View/Swiss-Pdb viewer (https://spdbv.vital-it.ch) and Molegro Molecular viewer (http://molexus.io/molegro-molecular-viewer) as described in previous studies ^24 30^. The coordinates of the BoNT/B lipid-binding loop (aa 1245-1252) present in PDB 2NM1 were inserted in PDBs files 4KBB and 6G5K. The sugar coordinates of GD1a were then merged with the PDB file 6G5K to reconstitute a trimolecular complex for SYT1. The preassembled complex GT1b/SYT1 and GT1b/SYT2 were docked on the synaptotagmin binding pocket according to the crystal coordinates of SYT1 (PDB 6G5K) and SYT2 (PDB 4KBB). All presented structures were of the BoNT/B1 subtype. The structures of the ceramide part of GD1a and cholesterol were retrieved from the CHARMM-GUI platform and added to the models to obtain a full system in a membrane context. Energy minimization of the complex with BoNT/B aa 1079-1290 was performed with the Polak-Ribière conjugate gradient algorithm, with the Bio+(CHARMM) force field in Hyperchem, typically with 3×10^5^ steps, and a root-mean-square (RMS) gradient of 0.01 kcal. Å^−1^.mol^-1^ as the convergence condition.

### Graphical representation of membrane-embedded complexes

In order to generate a schematic representation of the membrane-embedded protein complexes, the tool “membrane builder” available on CHARMM-GUI was used to generate a patch consisting of 128-DPPC (1,2-dipalmitoylphosphatidylcholine) molecules in a lipid-bilayer topology. Then, the minimized models were inserted according to the orientation proposed by the PPM Web Server. The snapshots were taken using Chimera software ^31^.

### Statistical analysis

Results are presented as mean ± SEM for immunofluorescence quantifications or mean ± SD for Langmuir monolayers and SPR analysis, of n independent experiments. Statistical analysis was performed using either Mann-Whitney U test or One-way ANOVA followed by Bonferroni post-hoc test for means comparisons. All statistical tests were performed using OriginPro 8.0.

## Results

### BoNT/B binds to SYT pre-assembled with GT1b

The JMD of SYT binds GT1b ^24^ via a consensus ganglioside-interaction motif that overlaps with the described BoNT/B-SYT interaction domain ^21^. We therefore addressed the question as to whether BoNT/B can bind to a SYT/GT1b complex. To investigate this point, we developed an SPR-based approach, consisting in capturing a peptide encompassing the JMD of SYT (pSYT1 32-57 or pSYT2 40-66, Supplemental Fig. 1) on a sensor chip, assembling a SYT/GT1b complex and then evaluating BoNT/B binding. GT1b diluted in running buffer interacted strongly with pSYT1 or pSYT2 immobilized on a sensor chip (Fig. 1a). The interaction was specific, as no binding occurred on a control pSYT9 peptide (Fig. 1a, Supplemental Fig. 1). Ganglioside binding to pSYT was detected at 10 nM GT1b (Supplemental Fig. 2a), a concentration far below its critical micellar concentration ^27^ and was dose-dependent (Supplemental Fig. 2b). Estimation of the ganglioside/peptide molar ratio indicated that a mean of 3 molecules of GT1b were bound per peptide (3.04 ± 0.4, n= 7 independent experiments ± SD using pSYT1 or pSYT2). This observation is compatible with ceramide-mediated multimeric self-assembly of gangliosides that occurs in lipid rafts^32^. The interaction of the JMD of SYT with GT1b was further corroborated using antibodies that specifically recognize the JMD of SYT. As shown in Supplemental Fig. 2c and d, GT1b bound to SYT, masks the recognition domain of anti-SYT JMD antibodies and inhibits their binding to SYT. Altogether, these results demonstrate that the ganglioside binding site of the JMD of SYT immobilized on a chip can stably capture GT1b. This experimental protocol mimics native conditions where SYT binds one GT1b molecule while being surrounded by other free GT1b molecules thus allowing BoNT/B to interact with SYT/GT1b and free GT1b using the SYT-binding pocket and GBS1 respectively.

**Fig. 1.**
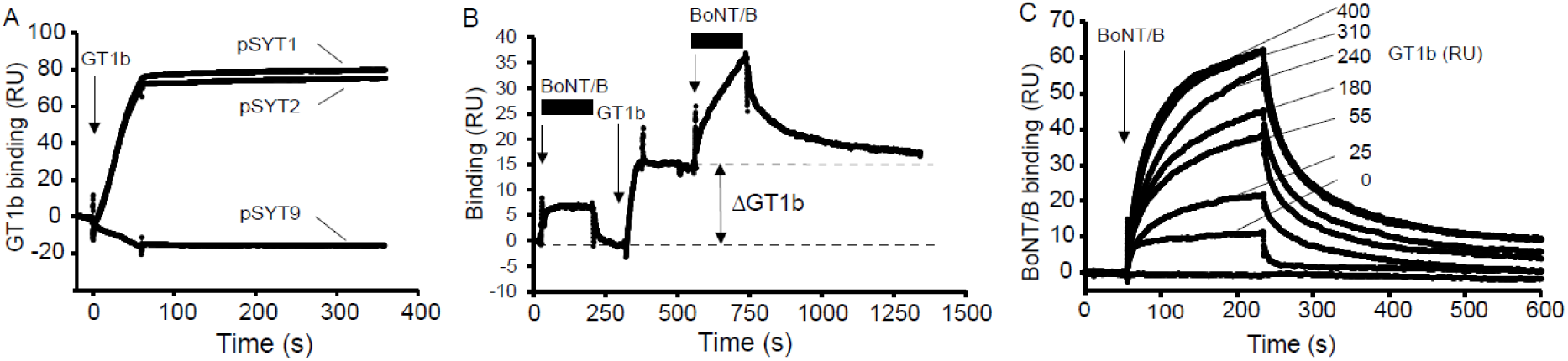
SPR based on-chip reconstitution of BoNT/B binding to SYT JMD pre-assembled with GT1b. **a** GT1b binding to pSYT1 and pSYT2. GT1b (200 nM) was injected for 1 min over immobilized pSYT1, pSYT2 and pSYT9 (240 RU) at 40 µl/min. Representative of >10 independent experiments. **b** GT1b bound to SYT induces an increment in BoNT/B binding signal. BoNT/B (30 nM) was injected (first arrow) onto pSYT1 (260 RU) showing a transient interaction that rapidly returns to baseline level (lower dashed line). GT1b (10 nM, second arrow) was then stably immobilized on pSYT1 (ΔGT1b) generating a new baseline (upper dashed line) and BoNT/B (30 nM, third arrow) was then injected again. Black bars highlight the BoNT/B injection phases. Representative of 6 independent experiments. **c** GT1b potentiation of BoNT/B binding to pSYT1 depends on the amount of GT1b bound to pSYT1. Sensorgrams resulting from the interaction of BoNT/B (30 nM) with pSYT1 (300 RU) pre-assembled with various amounts of GT1b (from 0 to 400 RU) were superposed. Representative of 3 independent experiments.

We then compared BoNT/B binding to SYT and SYT/GT1b complex. In the absence of GT1b, BoNT/B binding yielded a transient SPR signal on pSYT1 (Fig. 1b first arrow), consistent with its reported low affinity ^24 19^. GT1b was then immobilized on pSYT1 (Fig. 1b second arrow) before a subsequent injection of BoNT/B on the SYT/GT1b complex (Fig. 1b last arrow). Compared to pSYT1 alone, GT1b interaction with pSYT enhanced BoNT/B binding during the association phase, with a slower dissociation kinetic. This increase in BoNT/B affinity due to GT1b association with pSYT1 is similar to the effect of GT1b measured in proteoliposomes containing full length SYT1 ^24^. The potentiation of BoNT/B binding by GT1b depended on the amount of gangliosides bound to SYT, reaching a plateau (Fig. 1c, Supplemental Fig. 3a). At saturating concentrations of GT1b, BoNT/B binding to pSYT1 increase to 580 % ± 43 % (n=3 independent experiments ± SD). To rule out the possibility that GT1b alone produces an enhancement of BoNT/B binding independently of SYT, we used a mutant SYT1 peptide (F_46_A) that is unable to bind BoNT/B, but still interacts with GT1b ^20^. As shown in Supplemental Fig. 3a, in contrast to pSYT1/GT1b complex, BoNT/B did not interact with pSYT1-F_46_A/GT1b complex in agreement with the low affinity of BoNT/B for gangliosides. BoNT/B binding does not induce GT1b dissociation from pSYT/GT1b complex, as anti-GT1b antibodies detected the same amount of GT1b before and after BoNT/B binding (Supplemental Fig. 3b, c). BoNT/B binding to pSYT2 was also measured when GT1b was bound to SYT2, with the difference that BoNT/B signals were higher on pSYT2 than on pSYT1 in the absence of ganglioside, in accordance with their relative affinity (Fig. 1, Supplemental Fig. 3d). As for SYT1, the presence of GT1b bound to SYT2, promoted BoNT/B interaction with SYT2, increasing binding affinity mainly by decreasing the dissociation rate of BoNT/B from pSYT2 (Supplemental Fig. 3d). Altogether, these results demonstrate that BoNT/B binds to a preassembled SYT/GT1b complex.

### Molecular modeling of BoNT/B bound to a SYT/GT1b complex

In order to obtain molecular insight into the interaction of BoNT/B with SYT1-GT1b and SYT2-GT1b complexes, with specific information on the fate of the LBL upon toxin binding, we developed a molecular modeling strategy that takes into account membrane topology. Using the initial coordinates of toxin-SYT (PDB 4KBB, 6G5K and 2NM1), we constructed a full system consisting of a SYT-GT1b complex, a GD1a-toxin complex and a cholesterol molecule. After several rounds of energy minimization, a stable complex was obtained for both systems including SYT1 (Fig. 2a) and SYT2 (Fig. 2b). The initial conditions had to be slightly adjusted to take into account the TM domain of SYT and the ceramide part of the GD1a ganglioside which were absent from the crystal structure ^21^. These conformational constraints respected the overall geometry of the membrane, except that a gap between GD1a and the TM domain of SYT was filled by a cholesterol molecule ^16^. BoNT/B interacts with free GD1a and GT1b associated with SYT (∼25 % and ∼ 15 % of the total energy respectively) but in both cases the major contribution of toxin binding was due to SYT (∼ 60% of the total energy) as shown in the pie chart in Fig. 2. A key feature of our models is the insertion of the LBL loop between GT1b and SYT (Fig. 2, Supplemental Fig 4b). The overall energy of interaction of BoNT/B-SYT complex is around −600 kJ/mol (−592 kJ/mole for SYT1 versus −664 kJ/mole for SYT2). It was previously noted that the presence of ganglioside and SYT in contact with BoNT/B1 do not change its overall structure ^21^. This is also the case in our modeling data when a SYT/GT1b complex is bound to the toxin (Supplemental Fig. 5).

**Fig. 2.**
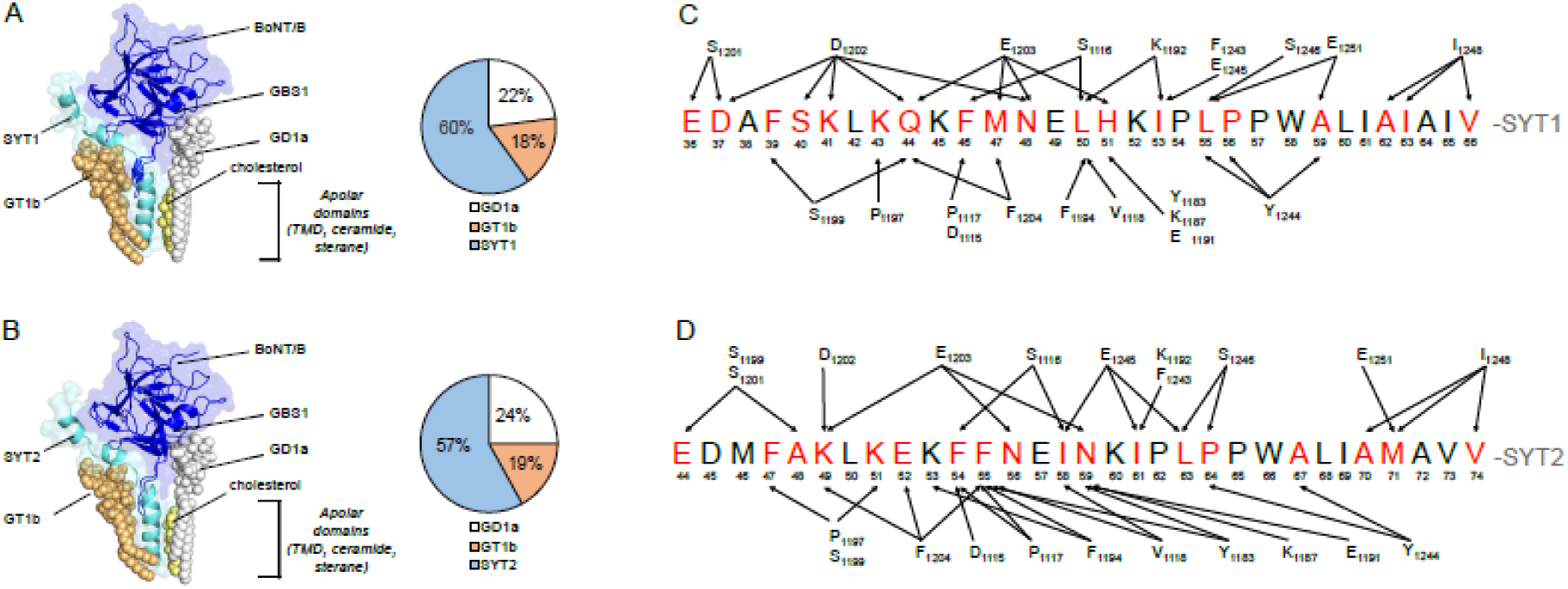
Overall structure of the energy-minimized complex between BoNT/B and its membrane ligands. **a** Model of SYT1-BoNT/B complex (SYT1 aa 34-72, BoNT/B HC aa 1079-1290). **b** Model of SYT2-BoNT/B complex (SYT2 aa 42-80, BoNT/B HC aa 1079-1290). The models are based on the initial superposition of a preformed synaptotagmin1/2-GT1b complex positioned in the SYT binding site of the toxin, also bound to GD1a, with a cholesterol molecule positioned between SYT-TM and GD1a ceramide. Both BoNT/B and SYT are represented as cartoons (dark and light blue respectively). The gangliosides and cholesterol are represented as spheres (GT1b: light orange, GD1a: white, cholesterol: light yellow). The apolar domains indicated in the models correspond to the sterane and isooctyl chains of cholesterol, the ceramide part of GD1a and GT1b, and the TMD of SYT. The pie charts indicate the relative distribution of the energies of interaction in the complex between BoNT/B and SYT1/2, GT1b, GD1a and between BoNT/B and SYT2, GT1b, GD1a. Note that cholesterol does not interact with the toxin, but with SYT TM and the ceramide part of GD1a. **c** and **d** depict mapping of the intermolecular interactions between BoNT/B and SYT. (**c**) SYT1 and (**d**) SYT2 residues interacting with the toxin are highlighted in red. Arrows indicate BoNT/B-SYT interaction points. A cut off <3.5 Å was used to select the illustrated residues.

#### BoNT/B-SYT interaction

The toxin was found to interact with a significant part of the extracellular regions of SYT1 (E_36_-W_58_) and SYT2 (E_44_-W_66_) via the SYT binding pocket, including loops Y_1183-_K_1188_, P_1197_-D_1202_ and K_1113_-P_1117_ (Supplemental Fig. 4a, Fig. 2c, d). The SYT helix, pre-conformed by GT1b, extends from E_36_ to H_51_ in SYT1 and E_44_ to N_59_ in SYT2, whereas a small distortion of the helix was observed in the central part of BoNT/B-SYT interface (Fig. 2a, b, Supplemental Fig. 4a). Almost all amino acids of BoNT/B that interact with SYT in crystal structures (6G5K, 2NP0, 4KBB, 2NM1) were found in contact with SYT and/or GT1b in our models (Supplemental Table 1). However, the conformational adjustment induced by the TM domain of SYT slightly turned the helix, generating a different interaction map with BoNT/B, compared to the BoNT/B-SYT binding interface determined by X-ray diffraction crystallography without membrane constraints. A detailed analysis of the contribution of amino acid residues of SYT and of the toxin revealed the evolution of the complex between the initial crystal conditions and the presented models (Supplemental Tables 1 and 2). SYT1-F_46_ and SYT2-F_54_, which initially interacted with GT1b in SYT/GT1b complexes ^24^, engage interactions with residues 1115-1117 of the toxin while retaining ganglioside association (Fig. 2c, d, Fig. 4, Supplemental Fig. 6). A superposition of the crystal structure (6G5K and 2NP0) with our models is shown in Fig. 3 for SYT1-F_46_ and M_47_ and its SYT2 counterparts that are described as key energetic hotspot residues in the crystal structure. In the case of SYT1 the aromatic ring of F_46_ (white) is replaced in the model by the apolar side chain of M_47_ (Fig. 3a, light blue). Consequently, most of the amino acid residues of the toxin that were in contact with F_46_ now interact with M_47_. In the case of SYT2 a similar substitution was evidenced between F_54_ in the crystal structure (white) and F_55_ in our model (Fig. 3b, light blue). Interestingly the aromatic ring of both residues is oriented in a similar way so that the pi-pi network involving residues Y_1183_, F_1194_ and F_1204_ of the toxin was still operative. These models uncover several additional SYT JMD residues compared to crystal data (E_36_, D_37_, S_40_, K_41_, Q_44,_ N_48,_ H_51_ in SYT1 and E_44_, A_48_, K_49_, E_52_, N_56_, N_59_ in SYT2) (Fig. 2c, d, Supplemental Table 2). Among them, SYT1-H_51_ and SYT2-N_59_ residues are facing the toxin and exhibit a high energy of interaction involving BoNT/B residues Y_1183_, K_1187_, E_1191_ and E_1203_ (Supplemental Fig. 7). In addition to the SYT/binding pocket, the BoNT/B LBL participates in the toxin-SYT complex by interacting tightly with apolar extramembrane (L_50_, I_53_, L_55_, P_56_) and membrane-embedded (A_59_, A_62_, I_63_, V_66_) residues of SYT1 (Fig. 2c, d, Supplemental Table 2a). Similar interactions were observed with homologous residues of SYT2 (Fig. 2c, d, Supplemental Table 2b).

**Fig. 3.**
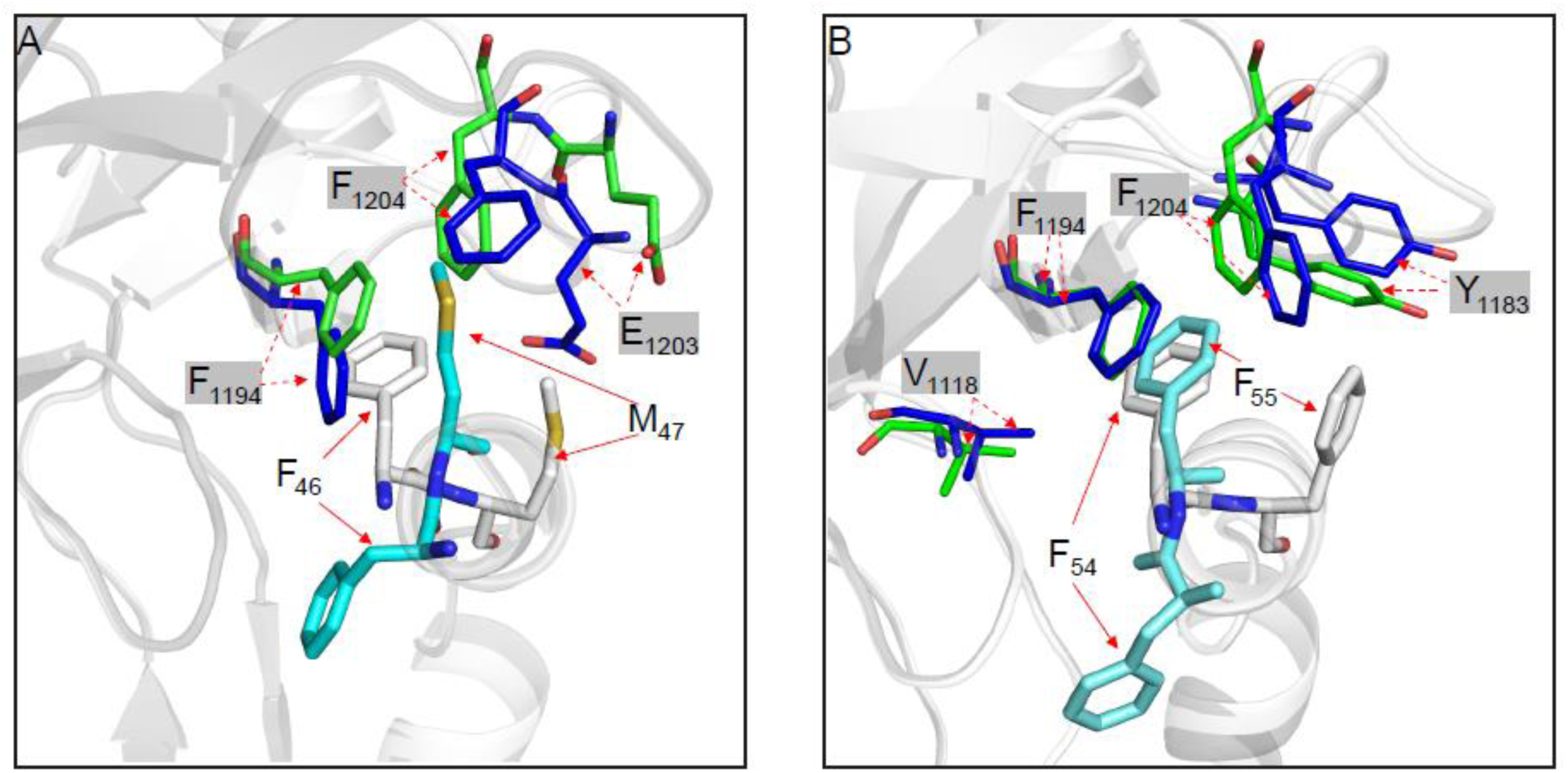
Close-up view of the molecular interface of SYT1-F_46-_M_47_ and SYT2-F_54_-F_55_. **a** Superposition of the SYT1-BoNT/B complex from PDB: 6G5K (BoNT/B in green and SYT1 in white) and the present model (BoNT/B in blue and SYT1 in light blue). **b** Superposition of the SYT2-BoNT/B complex from PDB 2NP0 (BoNT/B in green and SYT2 in white) and the present model (BoNT/B in blue and SYT2 in light blue). Interacting BoNT/B residues are shadowed in grey. Note that the position of BoNT/B residues are conserved between the proposed models and the corresponding crystal structures while their relative partners shift from F_46_ to M_47_ in SYT1 and F_54_ to F_55_ in SYT2.

#### SYT-GT1b interaction

GT1b imposes an angle of about 45° between the JMD of SYT and the membrane (Fig. 2a, b). The mapping of the molecular interactions between GT1b and either SYT1 (Fig. 4a) or SYT2 (Fig. 4b) and the toxin revealed that the binding involved the ceramide part of GT1b and the four terminal sugars and sialic acids (Glc1 and Gal2 are not involved in binding). The overall binding energy between SYT and GT1b was conserved upon interaction with BoNT/B (Supplemental Table 2). Interestingly, we noted a rearrangement involving F_46_ in SYT1 and its homologous F_54_ in SYT2 that reinforced the interaction with GT1b upon toxin binding (+32% and +90% for SYT1 and SYT2 respectively), involving Gal4, Sia5 and Sia6 (Fig. 4, Supplemental Table 2, Supplemental Fig. 8). In contrast, SYT1-K_52_, I_53_, L_55_, W_58_ and SYT2-K_60_, I_61_, W_66_ that were interacting initially with the Sia6 and Sia7 of GT1b in the preassembled SYT/GT1b complex, lose energy upon toxin binding (−92% and −47% respectively, Supplemental Table 2, Supplemental Fig. 8), suggesting also a molecular rearrangement in this region.

**Fig. 4.**
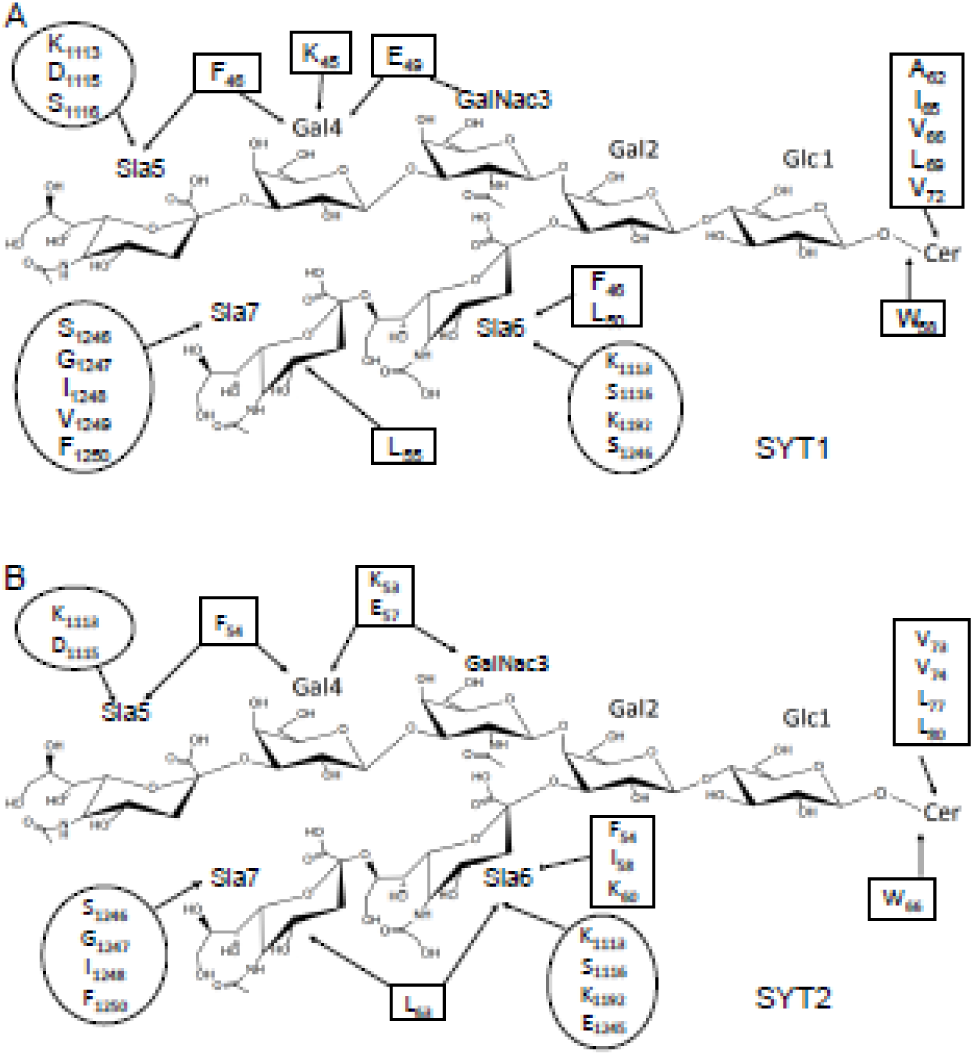
Schematic overview of intermolecular interactions between GT1b-BoNT/B and GT1b-SYT. The amino acids of SYT1 (**a**) and SYT2 (**b**) are boxed while those of the toxin are circled. A cut off <3.5 Å was used to select the residues indicated in the figure. Glc = glucose, Gal= galactose, Gal-Nac= N-acetylgalactosamine, Sia= sialic acid, Cer = ceramide. Only residues with energy ≥ 3 kJ/mol are listed

#### BoNT/B-GT1b and GD1a interactions

In the minimized complexes, both the LBL and the beta hairpin loop K_1113_-P_1117_ that interact with SYT, also bind to sialic acids of GT1b bound to SYT (Fig. 4, Supplemental Table 1b, Supplemental Fig. 9b). The LBL binds to Sia 7 whereas the K_1113_-P_1117_ loop binds to Sia 5 and Sia 6 (Fig. 4). Remarkably, as shown in Supplemental Fig. 9b, the BoNT/B loop K_1113_-P_1117_ corresponds to conserved β-hairpin loops E_1114_-V_1117_, A_1126_-R_1129_ and K_1143_-D_1147_ of BoNT/D, BoNT/C and tetanus toxin respectively which contribute to the sialic acid binding site ^33 12 13 34^. Our model suggests that the BoNT/B-protein binding pocket has an evolutionarily conserved ability to bind sialic acids that are brought by the SYT-associated ganglioside in the case of BoNT/B. Concerning the canonical ganglioside binding site, the BoNT/B residues interacting with the sugar part of GD1a were globally conserved after minimization compared to structural data (Supplemental Table 3).

Finally, it is worth noting that, in the minimized complex, cholesterol interacts with the TM domain of SYT, occupying a space created by the addition of the ceramide part of GD1a. Cholesterol increases the stability of the complex through a set of London forces with the ceramide part of GD1a and the TMD domain of SYT (Supplemental Table 4). These data raise the interesting notion that cholesterol could play an active role in the initial steps of toxin binding to lipid rafts in agreement with a previous description of SYT/cholesterol interactions ^16^.

Altogether these results suggest that BoNT/B interacts with the JMD and TMD domains of SYT along with two ganglioside molecules, one associated with SYT and the other with the ganglioside binding pocket of the toxin, with interconnection of the different intramembrane domains.

### The lipid binding loop of BoNT/B binds to GT1b

Our molecular modeling data suggest that the LBL could physically interact with SYT and GT1b. In order to corroborate LBL binding to GT1b we used the Langmuir monolayer method and recorded surface pressure changes in GT1b monolayers induced by a water-soluble BoNT/B peptide p1242-1256 encompassing the LBL (Supplemental Fig. 1). As shown in Fig. 5a, b, injection of p1242-1256 underneath a monolayer of GT1b yielded an increase in surface pressure (12.43 ± 1.14 mN/m, n=3), whereas limited interaction was found with lyso-LacCer (5.2 ± 0.72 mN/m, n=3), a lipid with an inverted conic shape resembling gangliosides, or sphingomyelin (1.97 ± 0.37 mN/m, n=3), a major sphingolipid component of lipid rafts. The absence of interaction with the ceramide domain of sphingomyelin indicates that the LBL of BoNT/B interacts preferentially with the sugar part of GT1b.

**Fig. 5.**
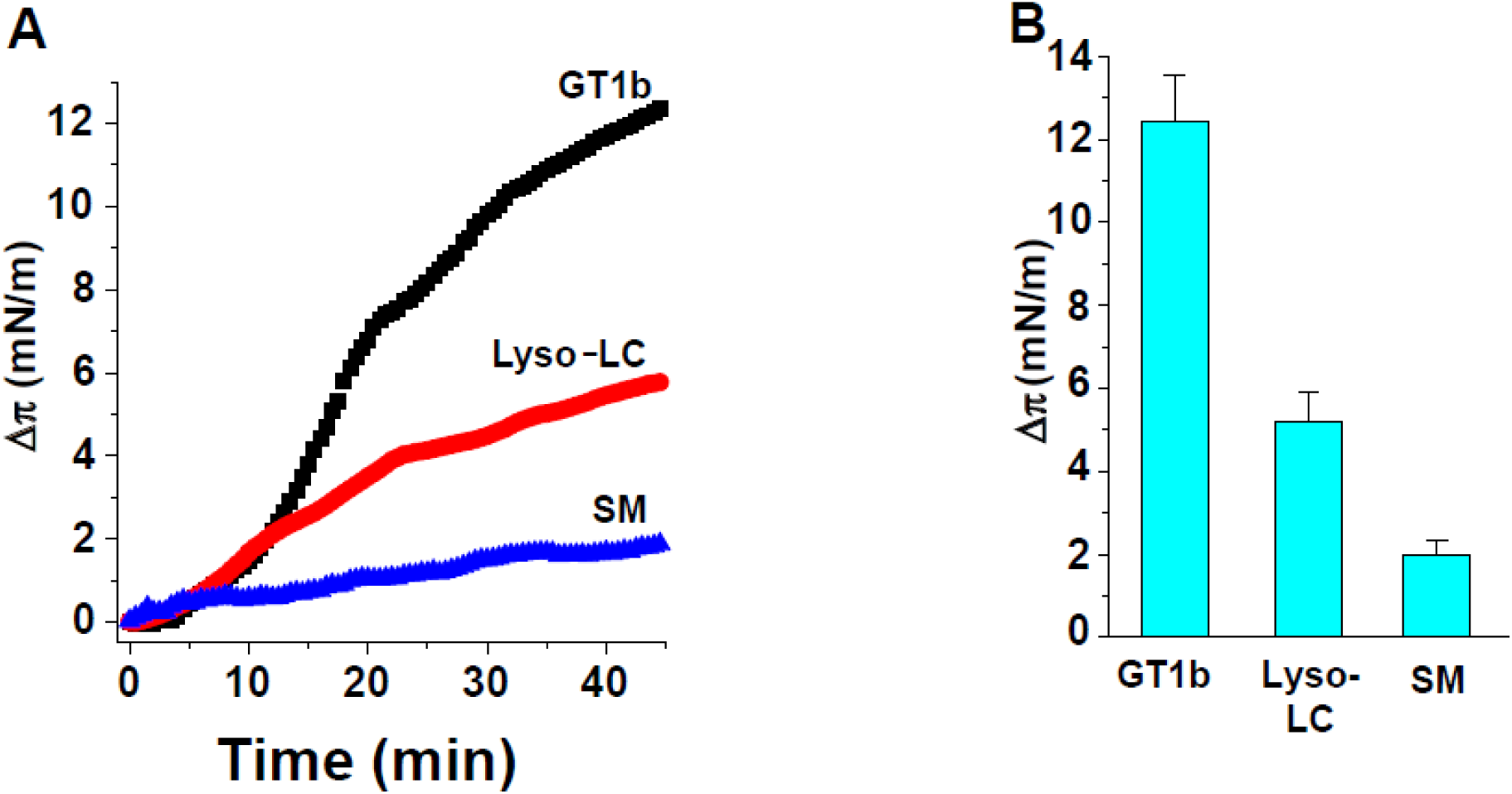
Measurement of BoNT/B apolar loop (LBL) interaction with GT1b using Langmuir monolayers. **a** Stable monolayers of GT1b, lyso-LacCer (Lyso-LC), and sphingomyelin (SM) were prepared at the air-water interface at an initial surface pressure of 15-20 mN.m^-1^. After equilibrium of the monolayer, the BoNT/B apolar loop (aa 1242-1256) was added at a final concentration of 10 µM. The data show the surface pressure increase Δπ induced by the loop as a function of time. The data are representative of three distinct experiments. **b** Histograms comparing the endpoints in a ± SD of three independent experiments.

### SYT1H_51_/K_52_ and SYT2-N_59_/K_60_ are key residues in BoNT/B binding

We have previously shown that the mutation SYT1-K_52_A abolished BoNT/B binding to PC12 neuroendocrine cells ^24^. Our present molecular modeling data predicts that, upon toxin binding, a molecular rearrangement occurs in the vicinity of SYT1-K_52_ and the corresponding SYT2-K_60_. We thus investigated whether the SYT2-K_60_ was also an important determinant in BoNT/B binding. Surface Plasmon Resonance analysis indicated that binding of the SYT2-G_40_-W_66_ peptide to GT1b-containing liposomes was drastically inhibited by mutating K_60_ to alanine (Supplemental Figs. 1 and 10), showing that SYT2-K_60_ residue is involved in the SYT2/GT1b interaction like SYT1-K_52_ ^24^. Immunofluorescence experiments showed that BoNT/B binding was severely impaired in PC12 cells expressing SYT2-K_60_A, compared to cells expressing SYT2-WT, with a decrease of 64 % (Supplemental Fig. 11a, c). A similar degree of inhibition (69%) was obtained using HEK 293 cells (Supplemental Fig. 11b, d). Although this decrease is less than that observed for SYT1-K_52_A ^24^, these results indicate that the SYT2-K_60_A like SYT1-K_52_, is an important determinant in BoNT/B binding.

According to our model, SYT1-H_51_ and SYT2-N_59_ residues adjacent to SYT1-K_52_ and SYT2-K_60_ respectively, exhibit a high energy of interaction with BoNT/B (Fig. 2c, d, Supplemental Fig. 7, Supplemental Table 2). To ascertain the functional involvement of SYT1-H_51_ in BoNT/B binding to SYT1, we mutated SYT1-H_51_ to a glycine residue and measured by immunofluorescence BoNT/B binding to either WT or SYT1 mutant transfected HEK 293 cells. The mutation SYT1-H_51_G induced a significant reduction (35%) in the binding of BoNT/B to the cell surface (IR BoNT/B of SYT1-WT: 1.00 ± 0.04 vs. IR BoNT/B of SYT1-H_51_G: 0.65 ± 0.04) while it did not affect expression levels of SYT1 (IR SYT of SYT1-WT: 1.00 ± 0.02 vs. IR SYT of SYT1-H_51_G: 1.01 ± 0.02) (Fig. 6). Interestingly, a variant of SYT2-N_59_ (SYT2-N_59_Q) occurs naturally in cats, which are known to be resistant to type B botulism ^35^, while expressing cleavable VAMP1, the predominant VAMP isoform in motor neurons ^6^ (Supplemental Table 5). Of note, a Q residue at position 59 was present in all Felidae sequences analyzed (Supplemental Table 5). We thus evaluated the potential impact on BoNT/B binding of this naturally occurring mutation. HEK 293 cells were transfected with either SYT2-WT or SYT2-N_59_Q and BoNT/B binding in the presence of GT1b was investigated by immunofluorescence. As shown in Fig. 6, the N_59_Q mutation induced a 50% decrease in BoNT/B binding (IR BoNT/B of SYT2-WT: 1.00 ± 0.03 vs. IR BoNT/B of SYT2-N_59_Q: 0.52 ± 0.02), while the expression of SYT2 remained unaffected (IR SYT of SYT2-WT: 1.00 ± 0.01 vs. IR SYT of SYT2-N_59_Q: 0.96 ± 0.01).

**Fig. 6.**
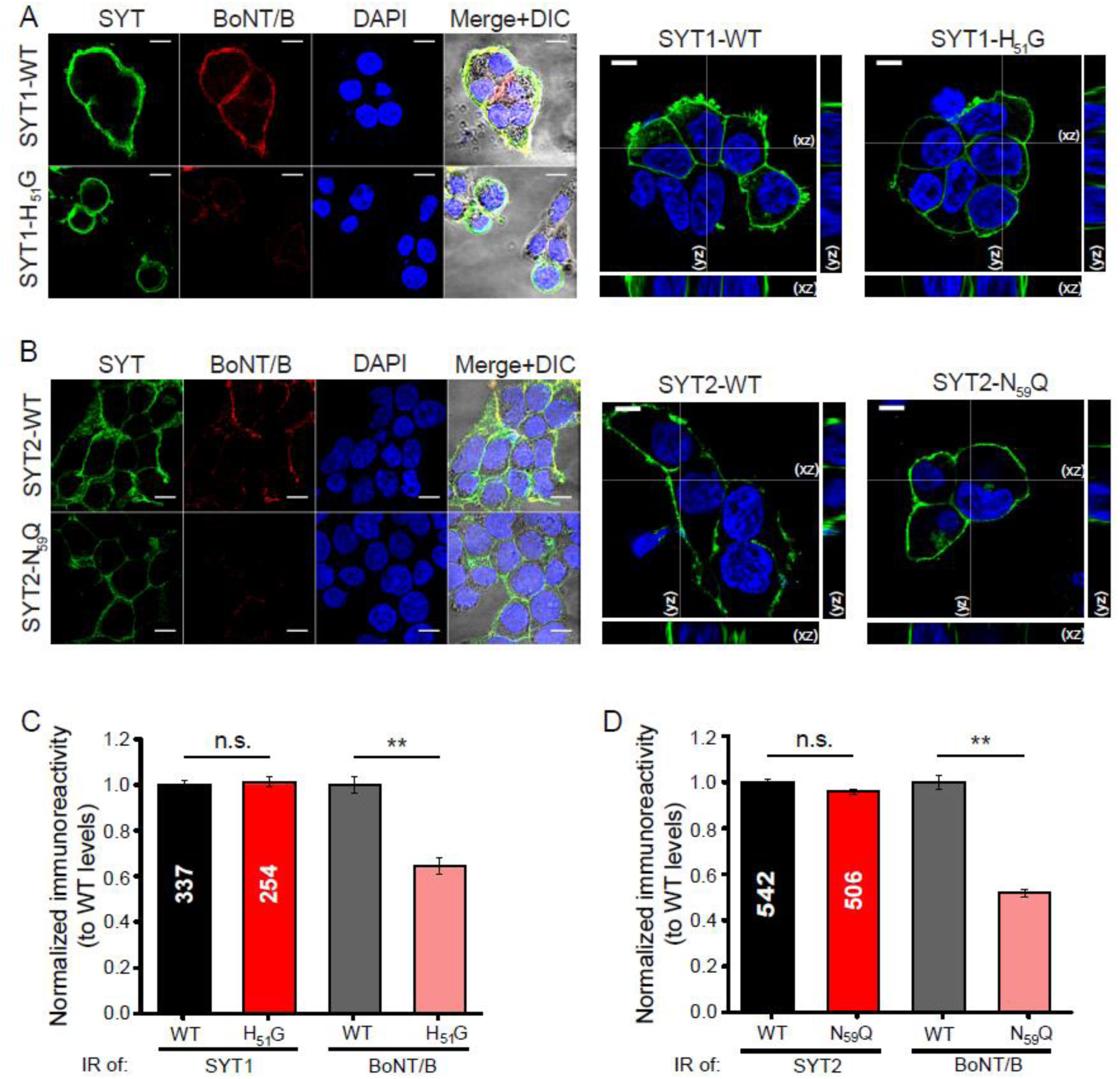
Mutations in the BoNT/B binding interface of SYTs decrease the binding of BoNT/B to HEK 293 cells. **a** Immunostaining of SYT1 (green) and BoNT/B (red) in HEK 293 cells transfected with either SYT1-WT (top) or H_51_G-SYT1 (bottom). DAPI signal is shown in blue, and the merge over DIC images indicated. Orthogonal projections of SYT1 labelling (green) in cells transfected with WT-SYT1 or SYT1-H_51_G (right). Scale bars, 10 µm. **b** Immunostaining of SYT2 (green) and BoNT/B (red) in HEK 293 cells transfected with either SYT2-WT (top panels) or SYT2-N_59_Q (bottom panels). DAPI signal is shown in blue, and the merge over DIC images indicated. Orthogonal projections of SYT2 labelling (green) in cells transfected with SYT2-WT or SYT2-N_59_Q (right). Scale bars, 10 µm. **c** Quantification of BoNT/B binding (grey and pink) and SYT1 expression (black and red) in cells expressing SYT1-WT or SYT1-H_51_G. The number of ROIs analyzed is indicated within each column. Normalized immunoreactivity data (IR) are expressed as mean ± SEM. Mann-Whitney U test was used for comparisons. **P < 0.01; n.s., non-significant. SYT1-WT IR to SYT1-H_51_G IR P=0.64; BoNT/B SYT1-WT to BoNT/B SYT1-H_51_G P=4.42 x 10^-12^. **d** Quantification of BoNT/B binding (grey and pink) and SYT2 expression (black and red) in cells expressing SYT2-WT or SYT2-N_59_Q. The number of ROIs analyzed is indicated within each column. Data are expressed as mean ± SEM. Mann-Whitney U test was used for comparisons. **P < 0.01; n.s., non-significant. SYT2-WT IR to SYT2-N_59_Q IR P=0.64; BoNT/B SYT2-WT to BoNT/B SYT2-N_59_Q P<0.001.

Taken together, these results indicate that both SYT1-H_51_K_52_ and SYT2-N_59_K_60_ homologous residues constitute a crucial doublet for BoNT/B binding, and are consistent with the proposed structure of the SYT/GT1b/GD1a/cholesterol/BoNT-B complex. They also provide an evolutionary explanation for the appearance of mutations (SYT2-N_59_Q) in animals with a diet at least partially based on carrion.

## Discussion

The current view of the BoNT/B binding determinants that anchor the distal tip of BoNT/B C-terminal domain to nerve terminals consists of two closed pockets interacting independently with SYT and gangliosides (GT1b or GD1a), and a lipid-binding loop thought to interact with the cell membrane via hydrophobic interactions. The central role of SYT in BoNT/B toxicity is supported by its relatively high affinity for the toxin, its synaptic localization conferring tissue specificity and by the observation that changes in potency among different BoNT/B subtypes are related to variability in the BoNT/B domain recognizing SYT but not in GBS1 ^36^.

Functional assays have unambiguously demonstrated that gangliosides are also necessary for BoNT/B intoxication and GT1b has a drastic synergistic effect on BoNT/B binding to SYT-containing membranes ^24 23 26 20^. However, the contribution of GT1b to neuronal membrane recognition by BoNT/B, is not totally understood. The affinity of the toxin for GT1b reconstituted in nanodiscs is weak (30-50 µM) ^14 15^ and not sufficient to measure detectable BoNT/B binding and VAMP2 cleavage in SYT knockout hippocampal neurons ^37^. Yet, both the canonical ganglioside binding site GBS1 and the LBL appear to participate in GT1b potentiation of BoNT/B binding to SYT, as inactivation of the GBS1 or the LBL abolish the synergistic effect of GT1b in vitro and cause a strong reduction of toxicity ^26 15^.

In a recent study we elucidated an important new mechanism underlying the role of gangliosides by demonstrating that GT1b actually binds to SYT JMD and induces the formation of an alpha-helix from an initially disordered domain ^24^. Intriguingly, GT1b overlaps critical residues defined by crystallographic and biochemical experiments in the SYT2-F_54_-I_58_ region, raising the question whether and how BoNT/B recognizes a preassembled GT1b/SYT complex and whether this complex dissociates upon BoNT/B binding ^24^.

Several experimental methods have been used to analyze the synergetic effect of GT1b on BoNT/B binding to SYT, including cultured cells and reconstituted systems (proteoliposomes, ELISA, mixed detergent micelles) ^24 23 26 38^. However, these approaches were not adapted to assessing whether during the GT1b potentiation effect, SYT is associated with GT1b. In the present study, we developed a SPR binding assay, with several GT1b molecules engaged in a complex with SYT, ensuring that BoNT/B could recognize SYT complexed to GT1b, along with free GT1b molecules that could also interact with the ganglioside-binding pocket of the toxin. Altogether, the SPR data strongly suggested that the SYT binding pocket of BoNT/B can accommodate SYT bound to GT1b.

We then used molecular modeling to assess how BoNT/B could recognize the SYT/GT1b complex. We docked the SYT/GT1b complex into the BoNT/B SYT-binding pocket, based on the co-crystal coordinates of BoNT/B associated with GD1a, together with the membrane-embedded domains of SYT and gangliosides. Cholesterol, which is known to interact with both the TM of SYT and the ceramide moiety of gangliosides^39^ was also included in the system. Flexible docking revealed that BoNT/B bound to GD1a can also interact with SYT1 or SYT2 bound to GT1b via the BoNT/B SYT-binding pocket described by structural data. The BoNT/B residues interacting either with SYT or GT1b are overall the same as described in the crystal structure, yet with additional interactions in the N-terminal domain of SYT. The SYT helix pre-conformed by GT1b extends from E_44_ to N_59_ and the BoNT/B-SYT interaction is mainly driven by apolar residues, including SYT2-F_47_, F_54_, F_55_ and I_58_ as previously described ^20 19^. In our present model, the membrane constraints induced by the overall polar and apolar ganglioside/SYT interactions, introduce an angle between SYT and the membrane plane, modifying the interaction map between BoNT/B and the SYT helix in comparison to structural data obtained with only the extracellular domain of SYT ^20 19 21^. It is of note that GT1b rescue of BoNT/B binding to several SYT mutants have revealed the crucial role of SYT2-F_54_ at the BoNT/B-SYT interface ^20 19 40^. Interestingly, our proposed models support this observation since they show that SYT2-F_54_ and its counterpart SYT1-F_46_ strongly interact with both GT1b and BoNT/B. Thus, molecular modeling, along with SPR binding experiments, are consistent with the view that the SYT/GT1b complex does not dissociate upon BoNT/B interaction and is recognized by the SYT binding pocket.

After the minimization process, the BoNT/B LBL was found to interact with the sialic acids of GT1b, the extracellular C-terminal part of the SYT JMD, as well as membrane-embedded SYT residues. This positioning of the LBL is allowed by the initial interaction of BoNT/B with the preformed SYT/GT1b complex, which determines the orientation of the toxin so that the LBL is directed toward the glycone part of GT1b and the TM domain of SYT. Consistent with our model, we determined experimentally that LBL directly binds to a monolayer of GT1b, but does not recognize sphingomyelin. This suggests that the LBL preferentially recognizes the polar headgroup of GT1b. The interaction between LBL and GT1b corroborates a report where LBL deletion decreased BoNT/B binding to nanodiscs containing GT1b ^15^. From a structural point of view, our findings suggest that BoNT/B LBL reinforces the interaction of BoNT/B with the SYT/GT1b receptor, explaining why its deletion dramatically reduces BoNT/B toxicity ^15^. Interestingly, BoNT/C, BoNT/D and tetanus toxin, also possess a lipid-binding loop that has been shown to bind sialic acids and it has been reported that the BoNT/C and BoNT/D LBL are structurally close to the corresponding SYT-binding site of BoNT/B ^13 34 12 41^. We propose that, for BoNT/B, C, D and tetanus toxin, a functional relationship exists between the presence of an LBL and the ability of these toxins to bind free or SYT-associated sialic acids outside of GBS1. As BoNT/G has a similar SYT-binding site to BoNT/B and an LBL, it would be interesting to investigate if this toxin also interacts with a SYT/GT1b complex ^40^. For BoNT/A and BoNT/E, the absence of LBL would be compensated by an interaction with glycosides covalently linked to their receptor, SV2 in this case ^1^.

In addition to the LBL, the model predicts that BoNT/B loop 1113-1117 also interacts with sialic acids of the GT1b-SYT complex. Interestingly and independently from GBS1, tetanus toxin, BoNT/C and BoNT/D use residues with a similar 3D-position to the BoNT/B 1113-1117 loop to mediate binding to sialic acids. As co-crystallization studies failed to detect the presence of sialyllactose in the BoNT/B SYT binding pocket ^42^, our model suggests that BoNT/B can bind sialic acid only when the glycoside is presented by SYT. The fact that BoNT/B could bind another ganglioside, in addition to that occupying GBS1, is compatible with the observation that trefoil recognition of carbohydrates is often multivalent ^43^. It has been noted that BoNT/B binds SYT using a pocket that is homologous to the sia-binding site of BoNT/C, BoNT/D and tetanus toxin ^12^, consistent with the view that the so-called sialic acid site of other BoNTs and the SYT binding site of BoNT/B may have a structurally related conserved function ^12^. Our model indicates that this pocket has conserved its ability to bind sialic acid associated with SYT in the case of BoNT/B. Thus, instead of the predominant view that BoNTs independently recognize a protein and a ganglioside or two gangliosides (BoNT/C) using two distinct pockets, our data support a new scenario in which a BoNT protein-binding pocket accepts a preformed protein/ganglioside complex. To our knowledge, this is the first description of recognition by a toxin of a protein/ganglioside complex.

Our model significantly extends our understanding of the BoNT/B-SYT binding interface and uncovers additional interacting residues in SYT. Among them SYT2-N_59_ was predicted to engage strong interactions with BoNT/B in particular with E_1191_, a key residue that modulates engineered BoNT/B activity for therapeutic purpose ^6^. Interesting, the natural variant SYT2-N_59_Q found in Felidae show a decrease in BoNT/B binding. To our knowledge type B botulism has never been reported in cats ^35^. These data may suggest that this natural variant could confer partial protection against type B botulism in animals feeding on carrion which can contain high amounts of BoNTs ^44^.

Our current findings thus highlight a new role for GT1b in BoNT/B binding by its direct interaction with SYT, explaining the poor affinity of BoNT/B for GT1b alone, but potently enhancing binding in the presence of SYT, particularly for SYT1. The very low affinity of BoNT/B for SYT1 has been suggested to be partially due to a steric clash with the toxin, involving SYT1-L_50_ ^20 45^. BoNT/B has been estimated to have at least 100-fold less affinity for SYT1 than SYT2 while GT1b reduces this difference 10-fold ^19 24^. Our present model predicts that SYT1-L_50_ binds GT1b and therefore a preassembled SYT/GT1b complex may facilitate BoNT/B binding to SYT1. Moreover our results are consistent with the fact that competitive neutralization of BoNT/B toxicity requires a SYT/ganglioside mixture rather SYT alone ^25^.

Our data revisit the dual receptor model ^46^ by uncovering an additional role for GT1b. We propose a new model in which, after the toxin is attracted and concentrated on the membrane by the negative charges of GT1b in lipid rafts ^47^, a preassembled and structured SYT/GT1b complex is accommodated in the SYT-binding pocket of BoNT/B, concomitantly to the binding of a ganglioside in the conserved ganglioside binding site GBS1. The LBL would then reinforce BoNT/B binding by interacting with GT1b associated with SYT. Accordingly, mutations in the GBS1, lipid binding loop or perturbation of GT1b/SYT interaction result in a loss of BoNT/B affinity and toxicity ^24 15 26^. After internalization, GT1b would participate in the toxin translocation process ^48^.

It has been suggested that simultaneous binding to SYT and gangliosides could impose a limited degree of freedom on BoNT/B orientation with respect to the membrane surface ^19 49^. In line with this notion, our data suggest that the intramembrane interactions between SYT and gangliosides could indeed immobilize both co-receptors at an appropriate distance, optimizing binding.

In summary, we present here a model of BoNT/B binding to neuronal membranes, that takes into account the specific topology of membrane receptors. BoNT/B has been successfully engineered to increase its affinity in a preclinical model ^6^. Our present findings could provide insights into the rational design of recombinant BoNTs for medical applications and for the development of inhibitors ^50 51 14^.

## Statements & Declarations

### Acknowledgments

We thank Raymond Miquelis for constructive discussions.

### Author contributions

CL, OEF and M Seagar conceived the study. CL and OEF supervised the entire project, the experimental design, data interpretation and manuscript preparation. CL and OEF analyzed and interpreted the data. JRF and CL performed immunofluorescence experiments, JRF acquired the corresponding images and performed treatment and analysis. FA performed molecular modelling and prepared with JRF molecular modelling figures. CL and GF performed SPR experiments. JF performed Langmuir monolayer experiments. FY and M Sangiardi performed expression plasmids preparation and preliminary expression tests. C.L. performed SPR experiments. C.L. wrote the original draft of the manuscript. CL, FA, JF, OEF and JRF prepared the figures. MRP was involved in discussion and data analysis. All authors edited and reviewed the manuscript.

### Ethics approval

Not applicable

### Consent to participate

All authors approved submission

### Consent to publish

All authors approved publication

### Data availability

Coordinates of structural Details of BoNT/B/synaptotagmin 1 & 2/gangliosides complexes are available upon request.

## Funding

Agence Nationale de la Recherche (ANR) (grant ANR-17-CE16-0022) for the postdoctoral financial support of JRF

Ministère des Armées (AID) and Aix-Marseille Université AMU for the PhD thesis of FO

## Abbreviations

BoNTs: Botulinum neurotoxins
HC: heavy chain
GBS1: ganglioside-binding site
SYT: synaptotagmin
LBL: lipid-binding loop
JMD: juxtamembrane domain
TMD: transmembrane domain
DMPC: 1,2-Dimyristoyl-sn-glycero-3-phosphocholine
DPPC: 1,2-Dipalmitoylphosphatidylcholine
IR: immunoreactivity
ROI: regions of interest
RMS: root-mean-square
Glc: glucose
Gal: galactose
Gal-Nac: N-acetylgalactosamine
Sia: sialic acid
Cer: ceramide.

