## Supplemental data for "Molecular landscape of BoNT/B bound to a membrane-inserted synaptotagmin/ganglioside complex"

**Electronic supplementary material**

| | |
|---|---|
| **Supplemental Fig.1** | Amino acid sequences of rat SYT and BoNT/B peptides used in this study |
| **Supplemental Fig. 2** | Characterization of GT1b interaction with the JMD of SYT |
| **Supplemental Fig. 3** | Characterization of BoNT/B binding to SYT/GT1b complex |
| **Supplemental Fig. 4** | BoNT/B-SYT molecular model with a membrane environment |

| | |
|---|---|
| **Supplemental Fig. 5** | Superposition of our current models and PDBs of BoNT/B-SYT complexes |
| **Supplemental Fig.6** | Close-up view of SYT1-F46 and SYT2-F54 interaction partners |
| **Supplemental Fig. 7** | Close-up view of the interaction interface of SYT1-$H_{51}$ and SYT2-$N_{59}$ with BoNT/B |
| **Supplemental Fig. 8** | Comparative energy profile of SYT/GT1b with or without toxin |
| **Supplemental Fig. 9** | A similarly positioned loop in BoNT/B, BoNT/D, BoNT/C and tetanus toxin binds sialyllactose |
| **Supplemental Fig. 10** | Effect of K60A mutation on GT1b binding to pSYT2 |
| **Supplemental Fig. 11** | Mutations in the $K_{60}$ residue of SYT2 inhibit the binding of BoNT/B to SYT2 expressing cells |
| **Supplemental Table 1** | Energy distribution of BoNT/B residues in contact with SYT and GT1b |
| **Supplemental Table 2** | Energy distribution of SYT residues in contact with BoNT/B and GT1b |
| **Supplemental Table 3** | Energy distribution of BoNT/B residues in contact with GD1a |
| **Supplemental Table 4** | Energy distribution of SYT residues in contact with cholesterol |
| **Supplemental Table 5** | Sequence alignment of SYT 1/2 and VAMP1 from different species |

| Peptides amino acid sequences | Peptide name |
|---|---|
| Biot-GEGKEDAFSKLKQKFMNELHKIPLPPW | pSYT1 (aa 32-58) |
| Biot-GESQEDMFAKLKDKFFNEINKIPLPPW | pSYT2 (aa 40-66) |
| Biot-GESQEDMFAKLKDKFFNEINAIPLPPW | pSYT2-$K_{60}A$ (aa 40-66) |
| Biot-GESQEDMFAKLKDKAANEINKIPLPPA | pSYT2 $F_{54}A$-$F_{55}A$-$W_{66}A$ (aa 40-66) |
| Biot-HDSCQDFIYHLRDRARPRLRDPDISVS | pSYT9 (aa 27-53) |
| RFYESGIVFEEYKDY | BoNT/B p1242-1256 (aa 1242-1256) |

**Supplemental Fig. 1** Amino acid sequences of rat SYT and BoNT/B peptides used in this study.

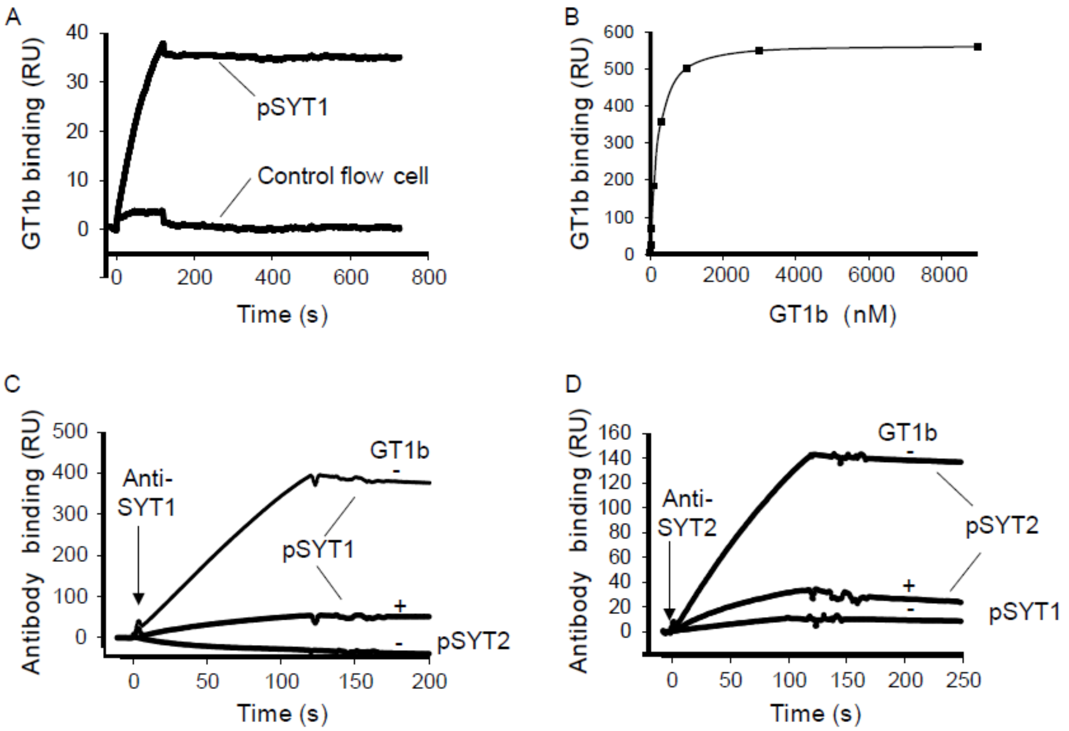


**Supplemental Fig. 2** Characterization of GT1b interaction with the JMD of SYT. **a** SPR measurement of the interaction of 10 nM GT1b with pSYT1. Traces represent the signal on the experimental flow cell functionalized with pSYT1 versus the signal on the control flow cell, indicating very low non-specific binding. **b** Dose-response curve of GT1b binding to pSYT1 (240 RU). GT1b (from 10 to 9000 nM) was diluted in HBS buffer and injected over pSYT1 versus control flow cell. **c** pSYT1 (510 RU) was immobilized on a sensor chip and probed with anti-SYT1 antibody (17 µg/ml) before or after GT1b binding (830 RU) to pSYT1. The specificity of the antibody is illustrated by an absence of interaction with pSYT2 and pSYT9 immobilized on control flow cell. Representative of 4 independent experiments. **d** pSYT2 (500 RU) was immobilized on a sensor chip and probed with anti-SYT2 antibody (10 µg/ml) before or after GT1b interaction (290 RU) with pSYT2 and pSYT9 immobilized on control flow cell. The specificity of the antibody is illustrated by an absence of interaction with pSYT1. Representative of 2 independent experiments.

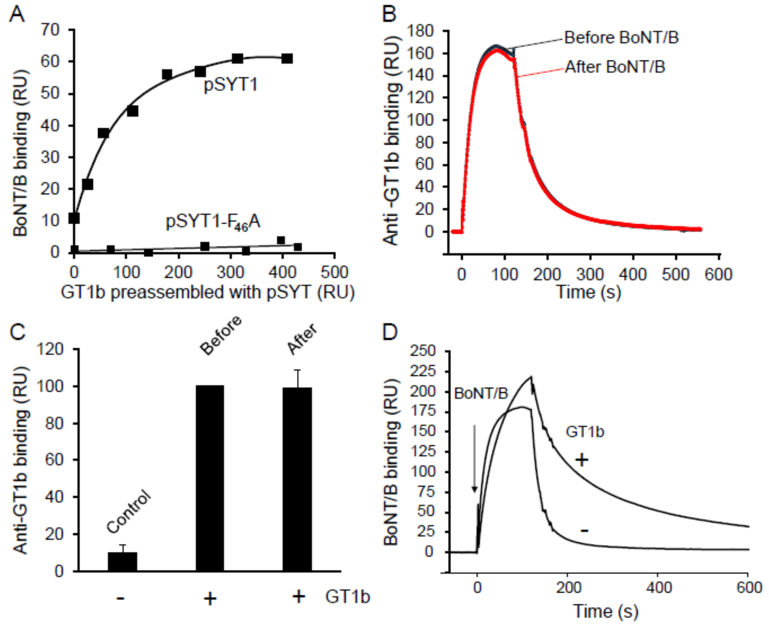


**Supplemental Fig. 3** Characterization of BoNT/B binding to SYT/GT1b complex. **a** Dose-response curve of the effect of GT1b on BoNT/B binding to immobilized pSYT1 or pSYT1-$F_{46}$A. Values taken 5 s before the end of injection from Fig. 1C were plotted (upper trace). Same experimental conditions were used to measure BoNT/B binding to pSYT1-$F_{46}$A (lower trace, representative of 3 independent experiments) **b** GT1b was captured over pSYT1 immobilized on a sensorchip. Anti-GT1b antibodies were then used to probe the level of GT1b bound to SYT before (black trace) and after injection (red trace) of BoNT/B (20 nM). Note that the antibody completely dissociates from GT1b after binding, allowing repetitive and comparative results. **c** Histograms of mean values ± SD obtained from 4 independent experiments conducted as in **b**. Anti-GT1b signals were normalized to GT1b signal bound to pSYT1 before toxin interaction. Control = background signal of anti-GT1b on pSYT1. Results are mean ± SD of 4 independent experiments. **d** GT1b potentiation effect of BoNT/B binding to pSYT2. Superimposed signals of BoNT/B (30nM) binding to pSYT2 immobilized (500 RU) on a sensor chip before and after GT1b binding to the peptide (400 RU).

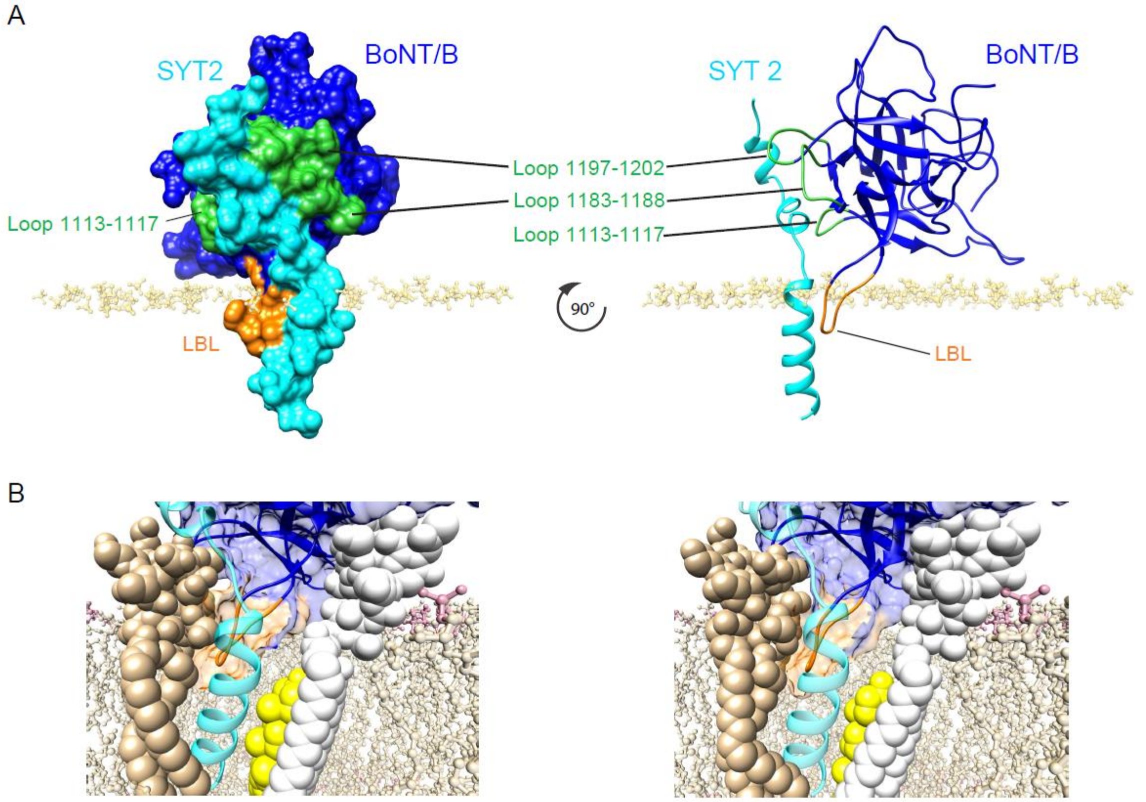


**Supplemental Fig. 4** BoNT/B-SYT representation with a membrane environment. **a** Surface representation (left) and cartoon representation (right) of SYT2 (light blue) complexed with BoNT/B (blue) showing their relative orientation to the plasma membrane. The loops of the BoNT/B-SYT2 binding pocket are depicted in green and the lipid binding loop (LBL: Loop 1245-1252) of BoNT/B is in orange. For the sake of visual clarity, only phosphate heads of DPPC (1,2-Dipalmitoylphosphatidylcholine) molecules are shown (pale yellow) and gangliosides omitted. Note the 90° rotation along the vertical axis of the complex (left and right panels). **b** Detail of the LBL penetration in a pseudo-realistic membrane context for SYT1 (left) and SYT2 (right) complexed with BoNT/B. SYTs are depicted in light blue, BoNT/B is in blue, GT1b is tan, GD1a is white, Cholesterol is yellow, and DPPC molecules are depicted in pale brown with the phosphate groups in pink. The LBL of both SYT1 and SYT2 is shown in orange.

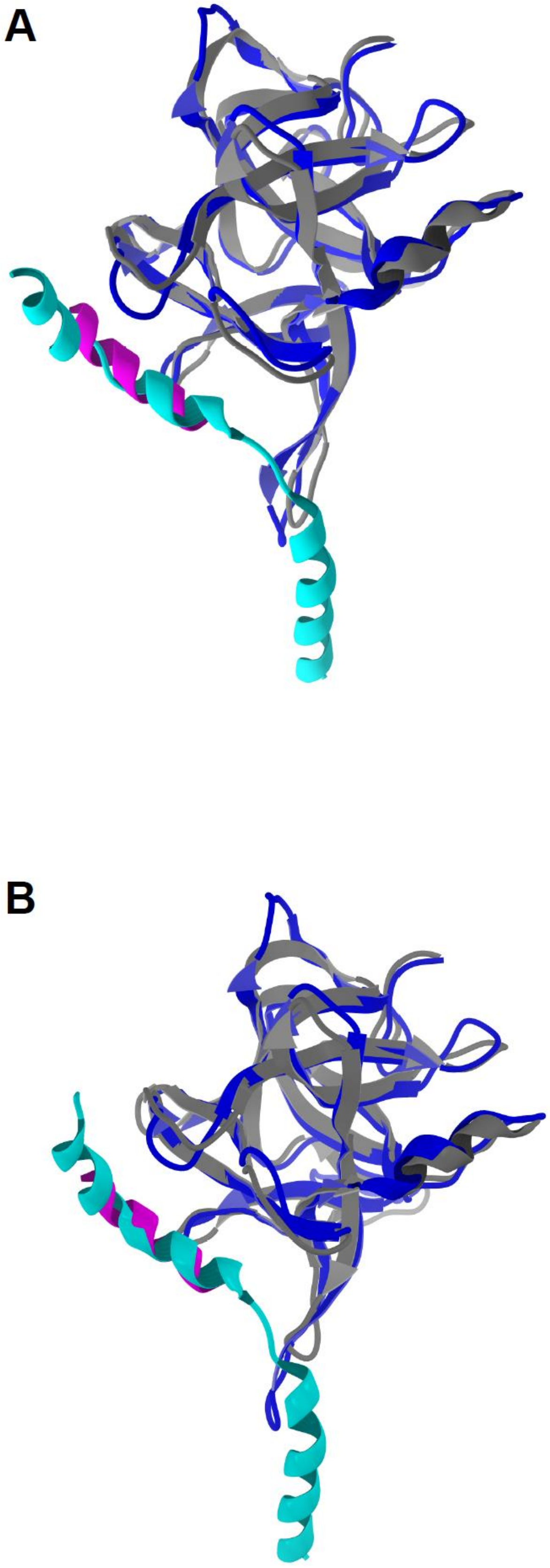


**Supplemental Fig. 5** Superposition of our current models and PDBs of BoNT/B-SYT complexes **a** SYT1: BoNT/B from our model (BoNT/B in dark blue and SYT1 in turquoise) and PDB 6G5K (BoNT/B in grey and SYT1 in pink) were superposed. **b** SYT2: BoNT/B from our model (BoNT/B in dark blue and SYT2 in turquoise) and PDB 4KBB (BoNT/B in grey and SYT2 in pink) were superposed. Gangliosides and cholesterol were omitted from this representation. Structures were superposed in Swiss-Pdb viewer and figures generated using Molegro.

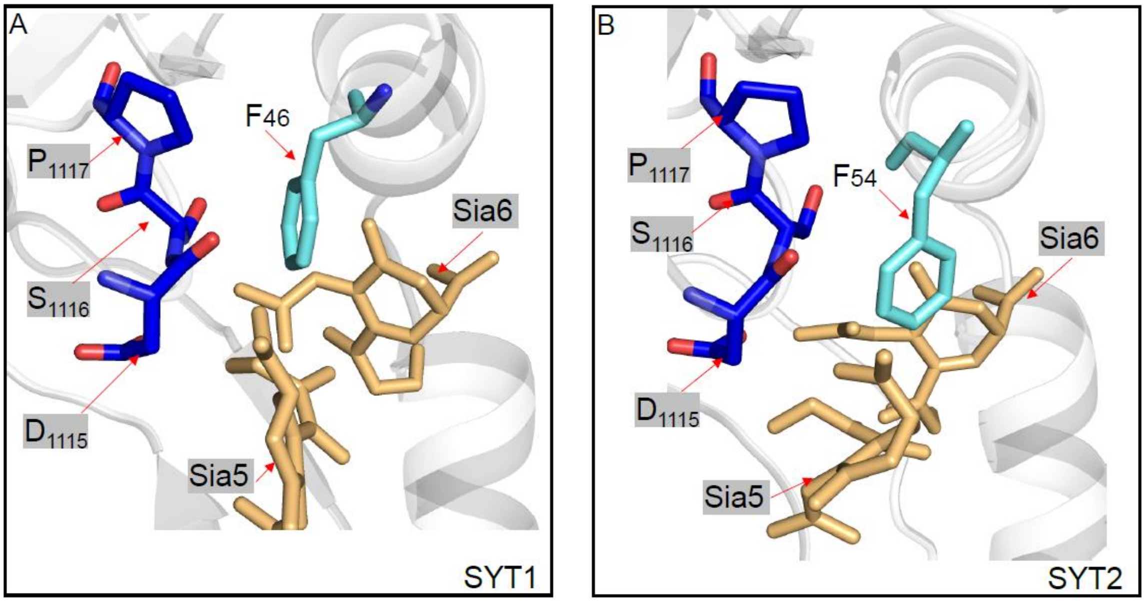


**Supplemental Fig. 6** Close-up view of SYT1-$F_{46}$ and SYT2-$F_{54}$ interaction partners. **a** SYT1-$F_{46}$ **b** SYT2-$F_{54}$. In both cases, the SYT aromatic residues (light blue stick) are interacting through a set of Van der Walls, CH-pi and OH-pi interactions with BoNT/B residues $D_{1115}$-$P_{1117}$ (blue stick) and sialic acid 5 and 6 of GT1b (yellow sticks). BoNT/B and SYT backbones appear as transparent light grey cartoons.

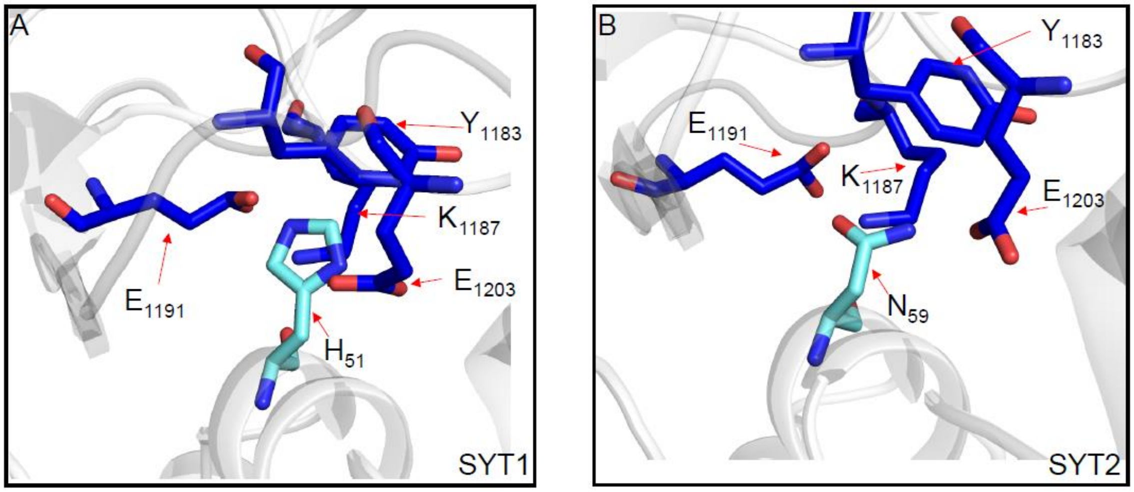


**Supplemental Fig. 7** Close-up view of the interaction interface of SYT1-$H_{51}$ and SYT2-$N_{59}$ with BoNT/B. Both residues SYT1-$H_{51}$ (**a**) and SYT2-$N_{59}$ (**b**) (light blue stick) interact with the same BoNT/B amino acids (blue sticks). BoNT/B and SYT appear as transparent light grey cartoons.

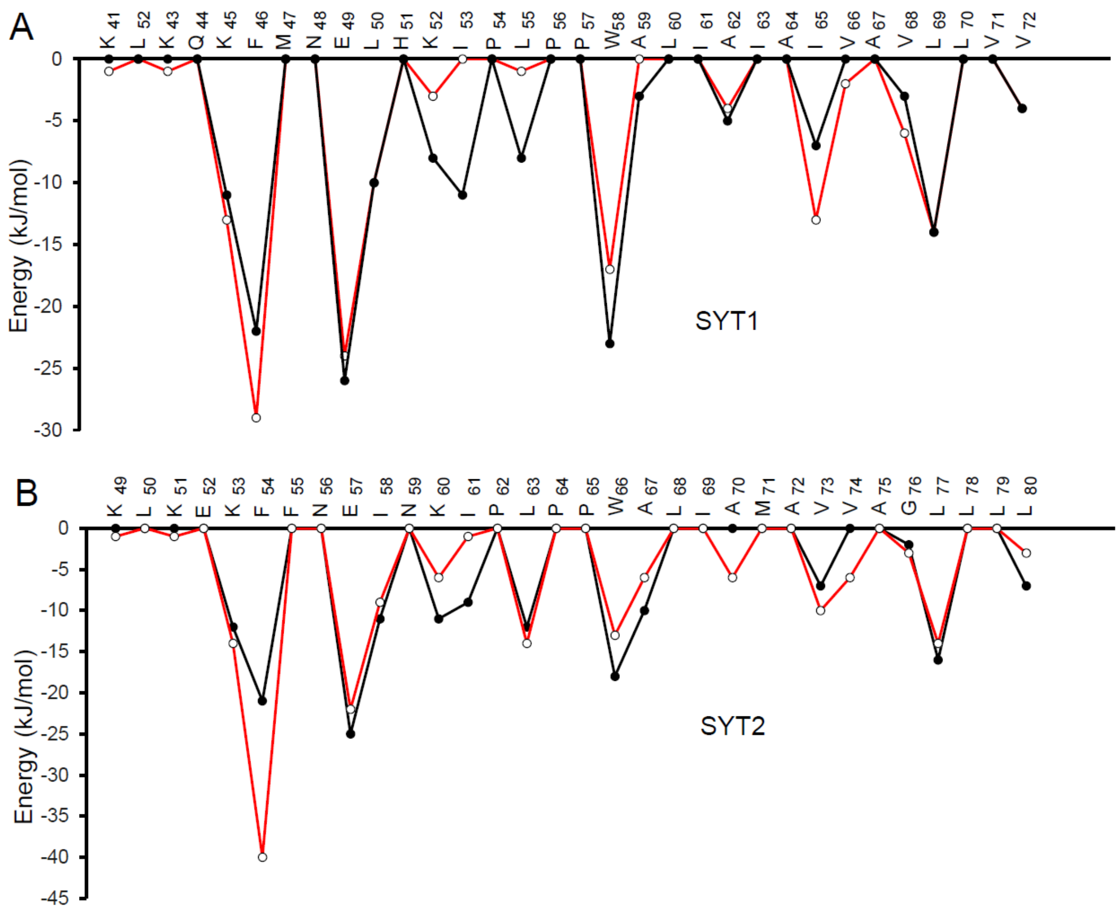


**Supplemental Fig. 8** Comparative energy profile of SYT/GT1b with or without toxin. Changes in SYT/GT1b energies (from Table S2) in the presence (red line) or absence (black line) of BoNT/B are depicted for SYT1 (**a**) and SYT2 (**b**).

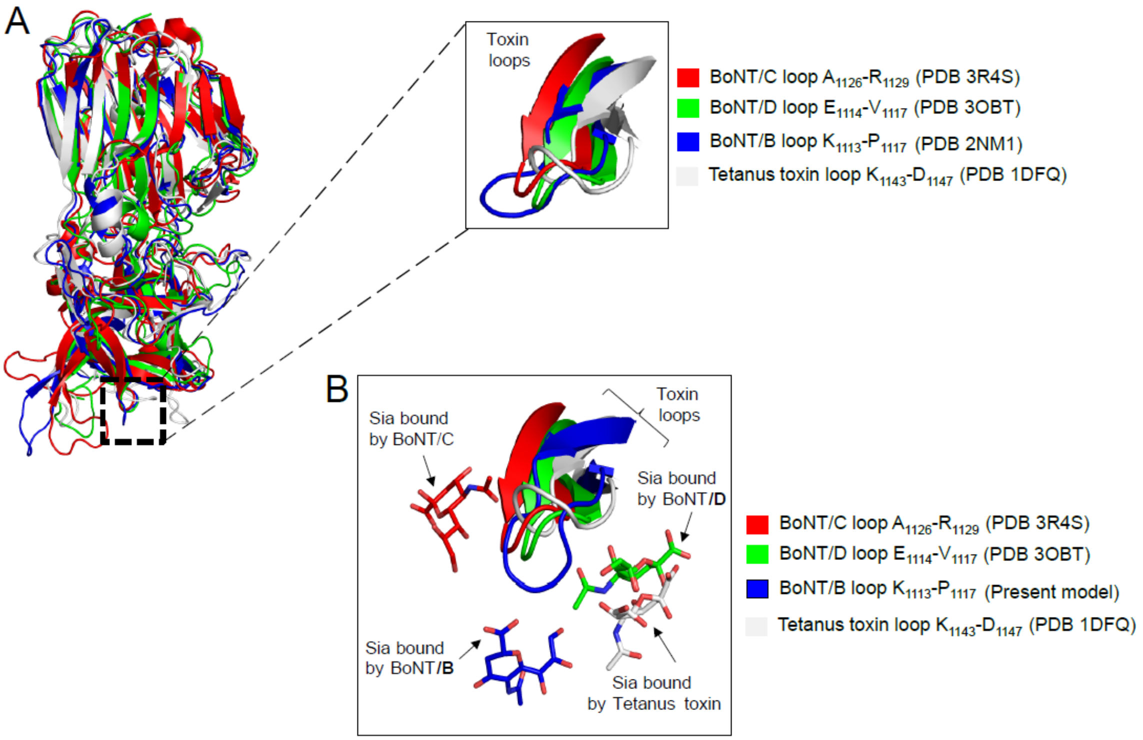


**Supplemental Fig. 9** A similarly positioned loop in BoNT/B, BoNT/D, BoNT/C and tetanus toxin binds sialyllactose. **a** (Left) Superposition of the crystallographic structure of BoNT/B (PDB 2NM1) with those of BoNT/C (red, PDB 3R4S), BoNT/D (green, PDB 3OBT) and tetanus toxin (white, PDB 1DFQ). (Right) The inset shows a close-up view of a β-hairpin loop overlay (BoNT/C loop $A_{1126}$-$R_{1129}$, BoNT/D loop $E_{1114}$-$V_{1117}$ Tetanus toxin loop $K_{1143}$-$D_{1147}$, BoNT/B loop $K_{1113}$-$P_{1117}$). **b** Same as (**a**) but the BoNT/B loop of PDB 2NM1 was replaced by the one issued from the present model and the interacting sialic acids (same code color as the corresponding toxins) from the used PDBs are depicted. The sialic acid interacting with the BoNT/B loop corresponds to the sia-5 of the SYT2 associated ganglioside in the present model.

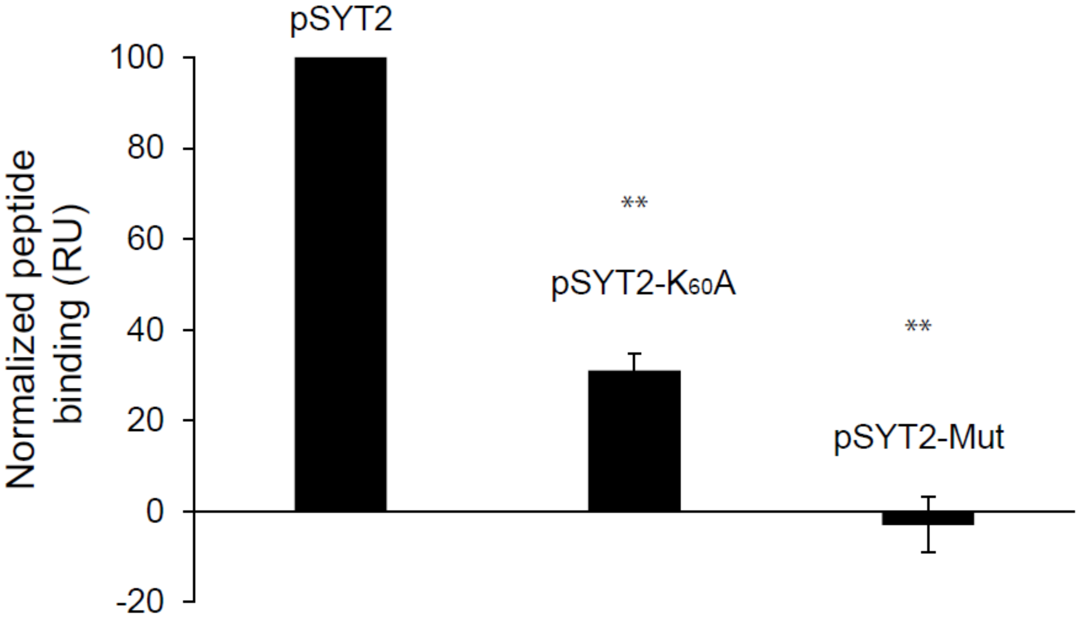


**Supplemental Fig. 10** Effect of $K_{60}A$ mutation on GT1b binding to pSYT2. Liposomes containing or not 8 % GT1b were immobilized on L1 chip and pSYT2, pSYT2-$K_{60}A$ and pSYT2-$F_{54}A$-$F_{55}A$-$W_{66}A$ (15 µM) injected. Compared to the signal with pSYT2, pSYT2-$K_{60}A$ induces a 71 % reduction of specific binding on GT1b whereas a triple mutation in the ganglioside binding domain abolished totally the signal. Results were normalized to pSYT2 binding and are mean of 4 independent experiments. One-way ANOVA followed by Bonferroni post-hoc test was used for means comparisons. **$P < 0.01$; pSYT2 WT vs pSYT2 $K_{60}A$: P= 5.15104 $x10^{-9}$; pSYT2 vs pSYT2-Mut P= $1.43928x10^{-10}$.

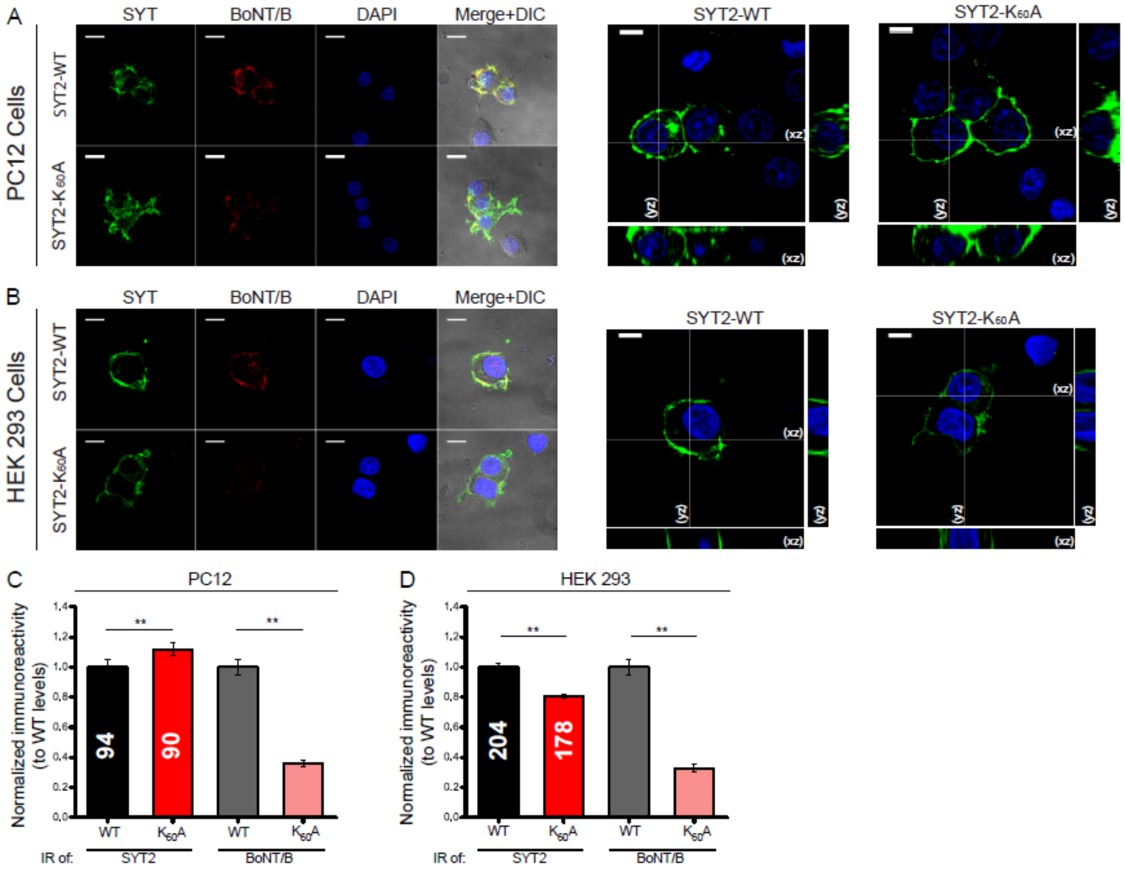


**Supplemental Fig. 11** Mutations in the $K_{60}$ residue of SYT2 inhibit the binding of BoNT/B to SYT2 expressing cells. **a** Immunostaining of SYT2 (green) and BoNT/B (red) in PC12 cells transfected with either SYT2-WT (top panels) or SYT2-$K_{60}$A- (bottom panels). DAPI signal is shown in blue, and the merge over DIC images indicated. Orthogonal projections of SYT2 labeling (green) in cells transfected with SYT2-WT or SYT2-$K_{60}$A (right). Scale bars, 10 µm. **b** Immunostaining of SYT2 (green) and BoNT/B (red) in HEK 293 cells transfected with either SYT2-WT (top panels) or SYT2-$K_{60}$A (bottom panels). Orthogonal projections of SYT2 labeling (green) in cells transfected with SYT2-WT or SYT2-$K_{60}$A (right). Scale bars, 10 µm. **c** Quantification of BoNT/B binding (grey and pink) and SYT2 immunoreactivity (black and red) in HEK 293 cells expressing SYT2-WT or SYT2-$K_{60}$A. The number of ROIs analyzed is indicated within each column. Normalized immunoreactivity (IR) data are expressed as mean ± SEM. Mann-Whitney U test was used for comparisons. $^{**}p < 0.01$; n.s., non-significant. SYT2-WT IR=1.00 ± 0.05; SYT2-$K_{60}$A IR=1.12 ± 0.04; BoNT/B IR on SYT2-WT=1.00 ± 0.05; BoNT/B IR on SYT2-$K_{60}$A=0.36 ± 0.02; SYT2-WT IR to SYT2-$K_{60}$A IR P=0.009; BoNT/B SYT2-WT to BoNT/B SYT2-$K_{60}$A P<0.001. **d** Quantification of BoNT/B binding (grey and pink) and SYT2 immunoreactivity (black and red) in PC12 cells expressing SYT2-WT or SYT2-$K_{60}$A. The number of ROIs analyzed is indicated within each column. Data are expressed as mean ± SEM. Mann-Whitney U test was used for comparisons. $^{**}P< 0.01$; n.s. non-significant. SYT2-WT IR=1.00 ± 0.02; SYT2-$K_{60}$A IR=0.81 ± 0.01; BoNT/B IR of SYT2-WT=1.00 ± 0.05; BoNT/B IR on SYT2-$K_{60}$A=0.31 ± 0.02; SYT2-WT IR to SYT2-$K_{60}$A IR $P=2.70 \times 10^{-9}$; BoNT/B SYT2-WT to BoNT/B SYT2-$K_{60}$A P<0.001.

**Supplemental Table 1** Energy distribution of BoNT/B residues in contact with SYT and GT1b. **a** Distribution of the energy of interaction of BoNT/B with SYT1 and SYT2. For comparison, interaction energies calculated from published structural data are listed. **b** Distribution of the energy of interaction of BoNT/B with GT1b in the SYT/GT1b complex. Residues corresponding to the LBL of BoNT/B are highlighted in grey in (**a**) and (**b**). The energies of interaction were obtained with Molegro molecular viewer. Only residues residues with energy ≥ 1 kJ/mol are listed.

A

| BoNT/B residues | Model SYT1 | Model SYT2 | PDB 6G5K SYT1 | PDB 2NP0 SYT2 | PDB 4KBB SYT2 | PDB 2NM1 SYT2 |
|---|---|---|---|---|---|---|
| | (KJ/mol) | | | | | |
| $K_{1113}$ | | | -8 | -12 | -9 | -13 |
| $D_{1115}$ | -6 | -5 | -5 | -4 | -9 | -6 |
| $S_{1116}$ | -11 | -19 | -6 | -8 | -6 | -8 |
| $P_{1117}$ | -4 | -9 | -17 | -16 | -17 | -16 |
| $V_{1118}$ | -1 | -5 | -7 | -6 | -6 | -6 |
| $W_{1178}$ | | | -3 | -3 | -3 | -4 |
| $Y_{1181}$ | -1 | | -6 | -2 | -1 | -5 |
| $T_{1182}$ | -1 | -1 | | | | |
| $Y_{1183}$ | -7 | -7 | -17 | -14 | -15 | -13 |
| $K_{1184}$ | | -1 | | | | |
| $Y_{1185}$ | | | -1 | | | |
| $F_{1186}$ | -1 | | | | | |
| $K_{1187}$ | -8 | -11 | | | | |
| $K_{1188}$ | | | -3 | | | |
| $E_{1189}$ | -1 | | | | | |
| $E_{1191}$ | -5 | -5 | -3 | -7 | -8 | -6 |
| $K_{1192}$ | -4 | -8 | -21 | -22 | -24 | -21 |
| $L_{1193}$ | | -4 | -4 | -3 | -3 | -4 |
| $F_{1194}$ | -23 | -24 | -35 | -31 | -31 | -34 |
| $L_{1195}$ | | | -1 | | | |
| $A_{1196}$ | | | -4 | -4 | -4 | -4 |
| $P_{1197}$ | -2 | -4 | -12 | -9 | -12 | -12 |
| $I_{1198}$ | -3 | -4 | | -1 | | |
| $S_{1199}$ | -9 | -14 | -8 | -7 | -6 | -12 |
| $D_{1200}$ | -1 | -3 | | | | -2 |
| $S_{1201}$ | -16 | -13 | -4 | -2 | -2 | -4 |
| $D_{1202}$ | -30 | -13 | -1 | | -1 | |
| $E_{1203}$ | -37 | -27 | -14 | -13 | -6 | -10 |
| $F_{1204}$ | -18 | -17 | -13 | -12 | -13 | -15 |
| $F_{1243}$ | -3 | -3 | | | | |
| $Y_{1244}$ | -21 | | | | | |
| $E_{1245}$ | -8 | -11 | -1 | | | |
| $S_{1246}$ | -6 | -14 | | | | |
| $G_{1247}$ | -1 | | | | | |
| $I_{1248}$ | -19 | -17 | | | | |
| $V_{1249}$ | -1 | | | | | |
| $E_{1251}$ | -9 | -3 | | | | |
| $K_{1254}$ | | -1 | | -1 | -1 | |
| $Y_{1256}$ | | | -3 | -4 | -5 | -2 |
| Total energy | -257 | -243 | -197 | -181 | -182 | -197 |

B

| BoNT/B residues | Model for SYT1 | Model for SYT2 |
|---|---|---|
| | (kJ/mol) | |
| $K_{1111}$ | -1 | -1 |
| $K_{1113}$ | -3 | -4 |
| $K_{1114}$ | -1 | -1 |
| $D_{1115}$ | -8 | -7 |
| $S_{1116}$ | -4 | -4 |
| $K_{1187}$ | -1 | -1 |
| $K_{1188}$ | | -1 |
| $K_{1192}$ | -3 | -3 |
| $R_{1242}$ | -1 | -1 |
| $E_{1245}$ | | -3 |
| $S_{1246}$ | -21 | -11 |
| $G_{1247}$ | -12 | -15 |
| $I_{1248}$ | -10 | -11 |
| $V_{1249}$ | -3 | -2 |
| $F_{1250}$ | -6 | -16 |
| $K_{1254}$ | -1 | -2 |
| $Y_{1256}$ | | -1 |
| Total energy | -75 | -84 |

**Supplemental Table 2.** Energy distribution of SYT residues in contact with BoNT/B and GT1b. **a** Left: Interaction energies of SYT1-BoNT/B in the present model and PDB 6G5K are listed. Grey indicates the SYT residues that interact with the LBL. Right: Interaction energies of SYT1-GT1b in the presence (this model) or absence [19] of BoNT/B. **b** Left: Interaction energies of SYT2-BoNT/B in the present model and PDBs 4KBB, 2NPO, 2NM1 are listed. Grey indicates the SYT residues that interact with the LBL. Right: Interaction energies of SYT2-GT1b in the presence (this model) or absence [19] of BoNT/B. Energies were calculated using the Molegro Molecular Viewer software and only residues with Energy ≥ 1 kJ/mol are listed. Conserved residues between the models and pdb files are highlighted in orange, residues present in PDB files and absent from our models are highlighted in blue and residues present only in our models are highlighted in green.

| A | SYT1 / BoNT/B Energy (kJ/mol) | | SYT1 / GT1b Energy (kJ/mol) | |
|---|---|---|---|---|
| SYT1 residues | Present model | PDB 6G5K | Present model | Model without BoNT/B |
| $E_{36}$ | -6 | | | |
| $D_{37}$ | -11 | | | |
| $A_{38}$ | | -2 | | |
| $F_{39}$ | -5 | -40 | | |
| $S_{40}$ | -11 | | | |
| $K_{41}$ | -14 | -1 | -1 | |
| $L_{42}$ | 0 | -13 | | |
| $K_{43}$ | -9 | -25 | -1 | |
| $Q_{44}$ | -20 | | | |
| $K_{45}$ | | | -13 | -11 |
| $F_{46}$ | -22 | -35 | -29 | -22 |
| $M_{47}$ | -36 | -5 | | |
| $N_{48}$ | -11 | | | |
| $E_{49}$ | | -18 | -24 | -26 |
| $L_{50}$ | -12 | -9 | -10 | -10 |
| $H_{51}$ | -37 | | | |
| $K_{52}$ | -2 | -2 | -3 | -8 |
| $I_{53}$ | -10 | | | -11 |
| $P_{54}$ | | | | |
| $L_{55}$ | -19 | | -1 | -8 |
| $P_{56}$ | -9 | | | |
| $P_{57}$ | | | | |
| $W_{58}$ | | | | -23 |
| $A_{59}$ | -7 | | | -3 |
| $L_{60}$ | | | | |
| $I_{61}$ | | | | |
| $A_{62}$ | -4 | | -4 | -5 |
| $I_{63}$ | -11 | | | |
| $A_{64}$ | | | | |
| $I_{65}$ | | | -13 | -7 |
| $V_{66}$ | -3 | | -12 | |
| $A_{67}$ | | | | |
| $V_{68}$ | | | -6 | -3 |
| $L_{69}$ | | | -14 | -14 |
| $L_{70}$ | | | | |
| $V_{71}$ | | | | |
| $V_{72}$ | | | -4 | -4 |
| Total energy | -259 | -150 | -135 | -155 |

| B | SYT2 / BoNT/B Energy (kJ/mol) | | | | SYT2 / GT1b Energy (kJ/mol) | |
|---|---|---|---|---|---|---|
| SYT2 residues | Present model | PDB 4KBB | PDB 2NP0 | PDB 2NM1 | Present model | Model without BoNT/B |
| $E_{44}$ | -21 | | | | | |
| $D_{45}$ | | | | | | |
| $M_{46}$ | | | | -4 | | |
| $F_{47}$ | -5 | -26 | -26 | -40 | | |
| $A_{48}$ | -8 | | | | | |
| $K_{49}$ | -32 | | -1 | -1 | -1 | |
| $L_{50}$ | | -11 | -12 | -12 | | |
| $K_{51}$ | -10 | -16 | -23 | -19 | -1 | |
| $E_{52}$ | -8 | | | | | |
| $K_{53}$ | | | -1 | 0 | -14 | -12 |
| $F_{54}$ | -25 | -36 | -34 | -34 | -40 | -21 |
| $F_{55}$ | -40 | -13 | -12 | -16 | | |
| $N_{56}$ | -8 | | | | | |
| $E_{57}$ | | -17 | -13 | -13 | -22 | -25 |
| $I_{58}$ | -13 | -15 | -13 | -12 | -9 | -11 |
| $N_{59}$ | -24 | | | | | |
| $K_{60}$ | | | | | -6 | -11 |
| $I_{61}$ | -8 | | | | -1 | -9 |
| $P_{62}$ | | | | | | |
| $L_{63}$ | -13 | | | | -14 | -12 |
| $P_{64}$ | -13 | | | | | |
| $P_{65}$ | | | | | | |
| $W_{66}$ | | | | | -13 | -18 |
| $A_{67}$ | -5 | | | | -6 | -10 |
| $L_{68}$ | | | | | | |
| $I_{69}$ | | | | | | |
| $A_{70}$ | -4 | | | | -6 | |
| $M_{71}$ | -10 | | | | | |
| $A_{72}$ | | | | | | |
| $V_{73}$ | | | | | -10 | -7 |
| $V_{74}$ | -4 | | | | -6 | |
| $A_{75}$ | | | | | | |
| $G_{76}$ | | | | | -3 | -2 |
| $L_{77}$ | | | | | -14 | -16 |
| $L_{78}$ | | | | | | |
| $L_{79}$ | | | | | | |
| $L_{80}$ | | | | | -3 | -7 |
| Total energy | -251 | -134 | -135 | -151 | -169 | -161 |

**Supplemental Table 3.** Energy distribution of BoNT/B residues in contact with GD1a. Interaction energies of BoNT/B and the sugar moiety of GD1a extracted from our model and PDB 4KBB of BoNT/B-GD1a complex.

| BoNT/B residues in interaction with GD1a | Model with SYT1 Energy | Model with SYT2 Energy | SYT2 (PDB 4KBB) Energy |
|---|---|---|---|
| | (kJ/mol) | | |
| $N_{1105}$ | -8 | -11 | -12 |
| $E_{1189}$ | | | -2 |
| $E_{1190}$ | **-7** | -6 | -11 |
| $I_{1240}$ | | -1 | -10 |
| $H_{1241}$ | -17 | -24 | -21 |
| $R_{1242}$ | -4 | -8 | -2 |
| $Y_{1253}$ | | | -2 |
| $S_{1260}$ | | | -3 |
| $W_{1262}$ | -15 | -10 | -41 |
| $Y_{1263}$ | -13 | -13 | -15 |
| $K_{1265}$ | | | -2 |
| $E_{1266}$ | -1 | -1 | |
| $R_{1269}$ | -2 | -2 | -1 |
| $N_{1273}$ | | | -1 |
| $K_{1275}$ | -22 | -23 | -23 |
| $L_{1276}$ | -4 | -5 | -9 |
| $G_{1277}$ | -1 | -4 | -7 |
| Total energy | -94 | -108 | -168 |

**Supplemental Table 4.** Energy distribution of SYT residues in contact with cholesterol. Energy of interaction between cholesterol and SYT1/2 residues in the modeled complexes.

| SYT1 residues in interaction with cholesterol | Energy (kJ/mol) | SYT2 residues in interaction with cholesterol | Energy (kJ/mol) |
|---|---|---|---|
| $L_{60}$ | -4 | $L_{68}$ | -4 |
| $I_{63}$ | -4 | $M_{71}$ | -9 |
| $A_{64}$ | -7 | $A_{72}$ | -9 |
| $A_{67}$ | -4 | $A_{75}$ | -6 |
| $V_{68}$ | -4 | $G_{76}$ | -4 |
| $V_{71}$ | -6 | $L_{79}$ | -14 |
| | | $L_{80}$ | -6 |
| Total energy | -29 | Total energy | -52 |

**Supplemental Table 5.** Sequence alignment of SYT 1/2 and VAMP1 from different species. SYT2 (aa $E_{44}$-$P_{65}$), SYT1 (aa $E_{36}$-$P_{56}$) and VAMP1 (aa $A_{71}$-$W_{91}$) sequences of mouse, cat, panther, cheetah and lynx are aligned and accession numbers of each sequence is listed.

| Species | | SYT2 ($E_{44}$-$P_{65}$) | Accession number | SYT1 ($E_{36}$-$P_{56}$) | Accession number | VAMP1 ($A_{71}$-$W_{91}$) | Accession number |
|---|---|---|---|---|---|---|---|
| Mouse | *(Mus. Musculus)* | EDMFAKLKEKFFNEINKIPLPP | NP_033333.2 | EDAFSKLKQKFMNELHKIPLPP | P46096 | ALQAGASQFESSAAKLKRKYW | NP_033522.1 |
| Cat | *(Felis catus)* | EDMFAKLKEKFFNEIQKIPLPP | XP_019677557.1 | EDAFSKLKEKFMNELHKIPLPP | XP_019690823.1 | ALQVGASQFESSAAKLKRKYW | XP_006933506.1 |
| Panther | *(Panthera pardus)* | EDMFAKLKEKFFNEIQKIPLPP | XP_019287308.1 | EDAFSKLKEKFMNELHKIPLPP | XP_019310737.1 | ALQVGASQFESSAAKLKRKYW | XP_019320298.1 |
| Cheetah | *(Acinonyx jubatus)* | EDMFAKLKEKFFNEIQKIPLPP | XP_026930522.1 | EDAFSKLKEKFMNELHKIPLPP | XP_026926989.1 | ALQVGASQFESSAAKLKRKYW | XP_014916890.1 |
| Lynx | *(Lynx canadensis)* | EDMFAKLKEKFFNEIQKIPLPP | XP_032447696.1 | EDAFSKLKEKFMNELHKIPLPP | XP_030179158.1 | ALQVGASQFESSAAKLKRKYW | XP_030177399.1 |
